# MicroRNA binding site variation is enriched in psychiatric disorders

**DOI:** 10.1101/2021.03.07.434312

**Authors:** Michael P. Geaghan, William R. Reay, Murray J. Cairns

**Affiliations:** The University of Newcastle, Callaghan, New South Wales, Australia; Centre for Brain and Mental Health Research, Hunter Medical Research Institute, New Lambton, New South Wales, Australia

## Abstract

Psychiatric disorders and other complex traits have a polygenic architecture, often associated with dozens or even hundreds of independent genomic loci. As each of these have a relatively small influence on the trait, the dissection of their biological components is a non-trivial task. For psychiatric disorders in particular, the majority of associated loci lie within non-coding regions of the genome, suggesting that most of the genetic risk for disease originates from the disruption of regulatory sequences. While previously exploration of the heritability of these sequences has focused on variants that modify DNA elements, those that alter *cis*-acting RNA sequences, such as miRNA binding sites, are also likely to have a significant impact in these disorders. MiRNA have already been shown to be dysregulated in these disorders through both genetic and environmental influence, so it is reasonable to suspect their target genes may also be affected by common variation. In this study, we investigated the distribution of miRNA binding site variants (MBSVs) predicted to alter miRNA binding affinity in psychiatric disorders and observed significant enrichment in schizophrenia, depression, bipolar disorder, and anorexia nervosa. We also observed significant enrichment of MBSVs in genes targeted by several miRNA families, including miR-335-5p, miR-21-5p/590-5p, miR-361-5p, and miR-557 in both schizophrenia and depression, and nominally significant enrichment of MBSV for miR-323b-3p in schizophrenia. We also identified a significant association between MBSVs in gene sets involved in regulation of the synapse and synaptic depression in schizophrenia. While these observations support the role of miRNA in the pathophysiology of psychiatric disorders, we also observed significant association of MBSVs in other complex traits suggesting that MBSVs are an important class of regulatory variants that have functional implications for many disorders.

## Introduction

Psychiatric disorders, including schizophrenia, major depressive disorder, bipolar disorder and autism spectrum disorders are prevalent to varying degrees in the general population (approximately 1% for schizophrenia, bipolar disorder and autism; approximately 35% for major depressive disorder), and are responsible for considerable morbidity and mortality. Existing treatments have a limited efficacy; for example, in the case of antipsychotics there is a 27% 1-year relapse rate (Leucht et al., 2012) and a range of detrimental side effects, such as cardiometabolic dysfunction (Lally & MacCabe, 2015) and other complications, which can reduce compliance. These drawbacks are partly due to a focus on symptoms, due in turn to a severe lack of understanding of the aetiological origins of these disorders.

In recent years, significant focus in the context of psychiatric disease has been given to a class of small, non-coding RNAs known as microRNAs (miRNAs) (M. Geaghan & Cairns, 2015). MiRNAs are important regulators of gene expression and translation. They recognise and bind to target mRNA molecules, typically within the 3’ UTR region *via* partial complementary binding to miRNA recognition elements (MREs). This complementary binding occurs primarily between the mRNA and the nucleotides 2-8 of the miRNA, referred to as the “seed region”. Active miRNA are associated with a protein complex known as the RNA-Induced Silencing Complex (RISC), which directs either silencing of protein translation or degradation of the mRNA molecule, thus resulting in post-transcriptional regulation of gene expression (Bartel, 2004). The short recognition sequence of mammalian miRNAs means that a single miRNA sequence is capable of targeting numerous genes, typically within the hundreds. Thus, miRNA-mRNA interaction networks are complex and differ from cell to cell and from tissue to tissue, dependent upon the local gene and miRNA expression profiles. While this complexity makes target prediction for miRNAs difficult, several algorithms have been developed to make credible bioinformatic predictions of the targets of miRNAs.

There is mounting evidence from both genetics and gene expression studies supporting a role for genetic and environmental alteration of miRNA function in psychiatric disease; indeed, the genetic locus containing the miR-137 host gene *MIR137HG* is host to one of the most significant common genetic associations with schizophrenia (Pardiñas et al., 2018; Schizophrenia Working Group of the Psychiatric Genomics Consortium, 2014). What has been less thoroughly studied is the possibility for an accumulation of genetic variants present within the MREs of genes involved in psychiatrically relevant networks. In the present study, we aimed to assess miRNA binding site variants (MBSVs) in nine psychiatric disorders with existing genome-wide association study (GWAS) summary data: schizophrenia (SCZ), bipolar disorder (BIP), major depressive disorder (MDD), autism spectrum disorders (ASD), post-traumatic stress disorder (PTSD), attention deficit hyperactivity disorder (ADHD), anorexia nervosa (AN), obsessive compulsive disorder (OCD) and Tourette’s syndrome (TS). We utilised the summary statistics from the latest GWAS for each disorder and the MiRNA Target Site database (dbMTS) (C. Li et al., 2020) to identify MBSVs that were significantly associated with each disorder. We further utilised a competitive gene set enrichment approach (de Leeuw et al., 2015) to identify pathways affected by MBSVs that were significantly associated with each disorder and performed a meta-analysis to assess whether any MBSV-affected pathways were associated with psychiatric disease in general. While these analyses revealed a significant enrichment of MBSVs in SCZ, BIP, MDD, ASD and AN, we also found evidence for aggregation of MBSVs within pathways relevant to synaptic function. These results suggest that MBSVs may contribute to the aetiology and/or pathophysiology of disorders such as schizophrenia and bipolar disorder, which suggests these may be valuable targets for future research into disease origins and novel treatments.

## Materials and Methods

### Genome-wide association study data

Genome-wide association study (GWAS) summary statistics were obtained from the Psychiatric Genomics Consortium website (https://www.med.unc.edu/pgc/download-results/) for the following nine psychiatric disorders: bipolar disorder (BIP), major depressive disorder (MDD), autism spectrum disorders (ASD), post-traumatic stress disorder (PTSD), attention deficit hyperactivity disorder (ADHD), anorexia nervosa (AN), obsessive compulsive disorder (OCD) and Tourette’s syndrome (TS). For schizophrenia (SCZ), the summary statistics from the more recent 2018 meta-analysis performed by the CLOZUK consortium (Pardiñas et al., 2018) were obtained (https://walters.psycm.cf.ac.uk/); similarly, for each PGC GWAS, the latest available summary statistics were used: BIP – 2018 (Stahl et al., 2019); MDD – 2019 (Howard et al., 2019); ASD – 2019 (Grove et al., 2019); PTSD – 2019 (Nievergelt et al., 2019); ADHD – 2018 (Demontis et al., 2019); AN – 2019 (Watson et al., 2019); OCD – 2018 (International Obsessive Compulsive Disorder Foundation Genetics Collaborative (IOCDF-GC) and OCD Collaborative Genetics Association Studies (OCGAS), 2018); TS – 2019 (Yu et al., 2019). A summary of the GWAS data is presented in Table 1.

**Table 1.**
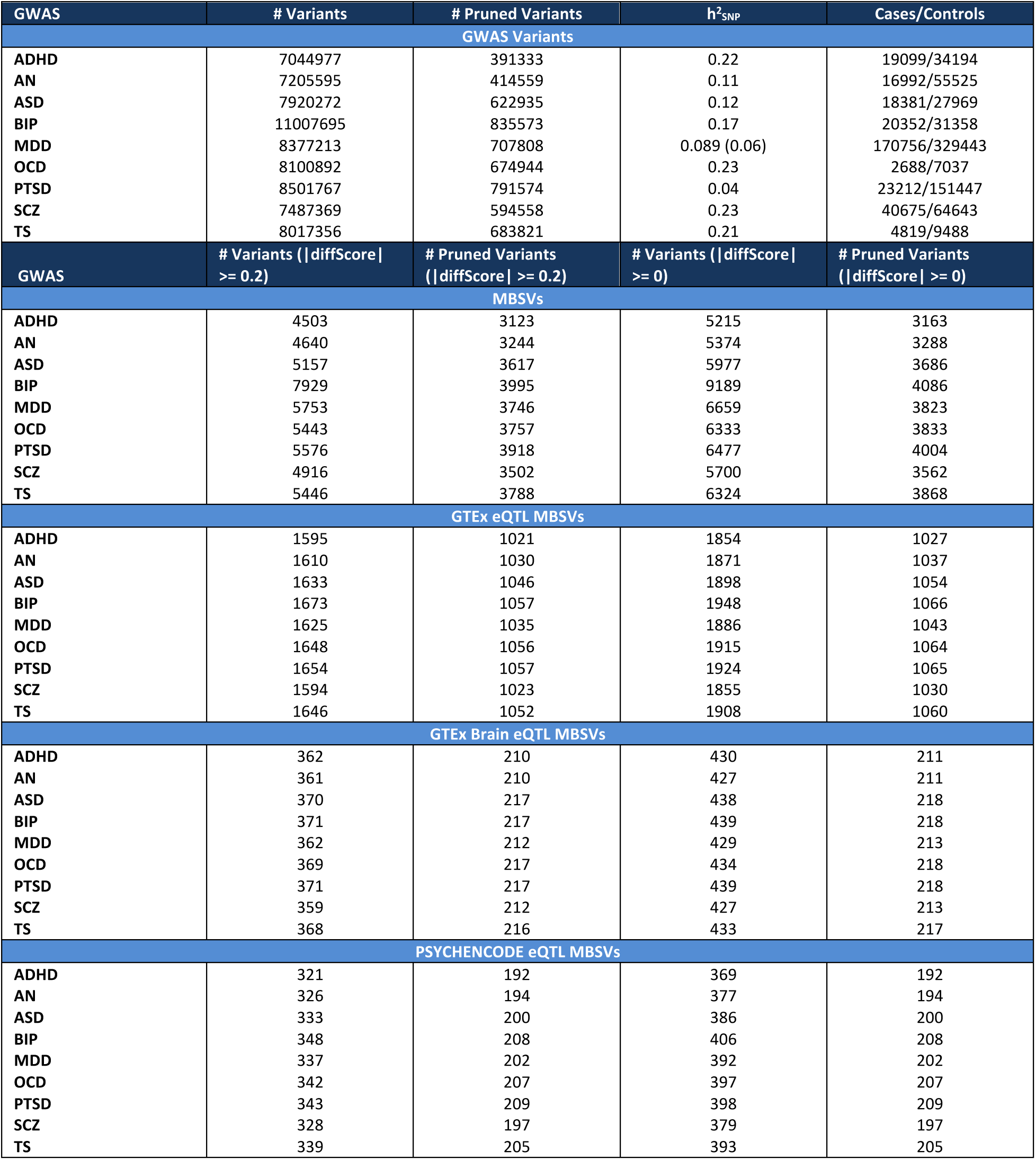
Summary of summary statistics and numbers of variants and MBSVs identified in each psychiatric GWAS.

### Identification of miRNA binding site variants

To identify all potential common miRNA binding site variants (MBSVs) associated with psychiatric disorders, we first took the union of all single nucleotide variants (SNVs) assessed in all GWAS. The chromosome, position, reference genome allele and alternate allele for each variant was identified. This information was then used to query the dbMTS database – an extension of the dbNSFP database containing all putative SNVs predicted to affect MREs based on three separate target prediction algorithms (TargetScan v7.0 (Lewis et al., 2005), miRanda (August 2010 release) (Enright et al., 2003; John et al., 2004) and RNAhybrid v2.1.1 (Rehmsmeier et al., 2004)).

#### Identification of brain-expressed genes and miRNAs

Both mRNA and miRNA brain expression data were obtained from the BrainSpan Atlas Developmental Transcriptome RNA sequencing dataset (v10) (J. A. Miller et al., 2014). This data was filtered to remove sequencing data from individuals less than 13 years of age, and represented samples from sixteen brain regions, including the dorsolateral prefrontal cortex (DFC), ventrolateral prefrontal cortex (VFC), anterior cingulate cortex (MFC), orbital frontal cortex (OFC), primary motor cortex (M1C), primary somatosensory cortex (S1C), posteroinferior parietal cortex (IPC), primary auditory cortex (A1C), posterior superior temporal cortex (STC), inferolateral temporal cortex (ITC), primary visual cortex (V1C), hippocampus (HIP), amygdaloid complex (AMY), striatum (STR), mediodorsal nucleus of thalamus (MD) and cerebellar cortex (CBC). The mRNA data was available as RPKM-normalised values. Genes were filtered for a minimum RPKM of 1 across at least 75% of samples. MiRNA data was available as raw read counts. These data were normalised to counts-per-million (CPM) by dividing the raw read counts by the total read count (in millions) of the respective samples. Then, the miRNAs were filtered for a minimum CPM of 10.30 (equivalent to a raw count of 10 in the smallest library) across 75% of samples. The miRNAs were further filtered for confidently-annotated miRNA species, as determined by the TargetScan database (v7.2). This includes miRNAs annotated by TargetScan as “highly conserved”, “conserved” and “poorly conserved but confidently annotated”, and excludes the “poorly conserved and possibly misannotated as a miRNA” category.

#### Variant filtering

Non-autosomal variants were filtered out prior to any analysis. Variants were further filtered by the expression of the affected miRNAs and mRNAs within the brain. Next, information on both the reference and alternate alleles for each variant was extracted from the dbMTS data. Alleles were considered to overlap a “true” MRE if both TargetScan and at least one of the other two algorithms annotated it for the same miRNA and gene. For each surviving allele, the TargetScan predictions were then used for all remaining analyses.

#### Calculation of variant scores

For each allele-miRNA-transcript combination, the dbMTS data contained a target prediction score calculated by TargetScan. For each gene, the best (i.e. most negative) score for each allele-miRNA combination across all transcripts was retained as a representative score for that allele-miRNA-gene interaction. Next, a representative “difference score” for each variant-gene interaction was calculated by subtracting the best reference allele score from the best alternate allele score. For the subset of variants that affected the 3’ UTRs of multiple genes, the best difference score was retained as the representative score for that variant. As such, each score represented the effect a variant had on the optimal miRNA binding conditions. A positive difference score represented a loss of miRNA binding affinity, and a negative score represented a gain, reflecting the negative scoring system of TargetScan. Furthermore, for many of the downstream analyses, “true” MBSVs were considered to be those with an absolute difference score ≥ 0.2, which equated to removing the lower 15% of scores.

### Annotation of eQTLs

The dbMTS data contained eQTL information for each variant obtained from the GTEx database v6 (Lonsdale et al., 2013), including each gene and respective tissue for which the variant was an eQTL. MBSVs were considered an eQTL if they affected the same gene that was annotated in the GTEx data. Furthermore, MBSVs were considered brain-eQTLs if they affected the same gene in any brain tissue. Additionally, variants were cross-referenced with brain-eQTLs identified by the PSYCHENCODE consortium (D. Wang et al., 2018). PSYCHENCODE eQTLs were filtered by an FDR < 0.05 and an FPKM > 0.1 in at least 150 samples.

### Annotation of CADD scores

We annotated all GWAS variants, including MBSVs for CADD (combined annotation dependent depletion) scores (Rentzsch et al., 2019). CADD scores were obtained directly from the CADD SNV database, using both the latest available version (v1.6) in addition to the original published version (v1.0) (Kircher et al., 2014). The original database was used to determine if CADD scores were biased for MBSVs due to the inclusion of miRNA target prediction score annotations in the generation of the scores from v1.1 onwards.

### Association of MBSVs with psychiatric disorders

To assess whether MBSVs were associated with psychiatric disorders, we first investigated the empirical cumulative distribution functions (ECDFs) of p-values associated with MBSVs in each disorder and compared these to those of all non-MBSV GWAS variants. MBSVs were also compared against all non-MBSV 3’ UTR variants. ECDFs for eQTL subsets of MBSVs were also calculated. Variants were first pruned using PLINK (v1.90) using the European subset of the 1000 Genomes Phase 3 cohort (N = 503), a window and step size of 50 and 5 nucleotides respectively, a minor allele frequency (MAF) threshold of 0.01 and an R^2^ threshold of 0.1. When pruning, MBSVs were prioritised. ECDFs were compared using the Kolmogorov-Smirnov (K-S) test. ECDFs of the absolute log effect sizes (i.e. log odds ratio or beta) were also calculated and compared. For each disorder, the proportion of MBSVs that were genome-wide significant (p < 5x10^-8^) and suggestive-significant (p < 1x10^-5^) was compared to the proportion of either all non-MBSV variants or non-MBSV 3’ UTR variants, using Fisher’s exact test to assess whether there was any enrichment or depletion of MBSVs. For both K-S and Fisher’s exact tests, the false discovery rate (FDR) across all tests for each disorder was calculated using the Benjamini-Hochberg method, and an FDR < 0.05 was considered significant.

We next aggregated MBSV p-values obtained from the respective GWAS summary statistics, in a manner analogous to traditional gene-set association analysis. In order to account for correlation between p-values for MBSVs in linkage disequilibrium (LD), we applied the aggregated Cauchy association test (ACAT) method (Liu et al., 2019). The advantage of this method is its computational simplicity and its low sensitivity to correlated p-values, particularly when the aggregated p-value is small. This method takes p-values *p*_*i*_ for *i* = (0, 1, ⋯, *k*) and optional non-negative weights *w*_*i*_ and returns an approximate p-value *p*_*ACAT*_ representing the aggregated association:

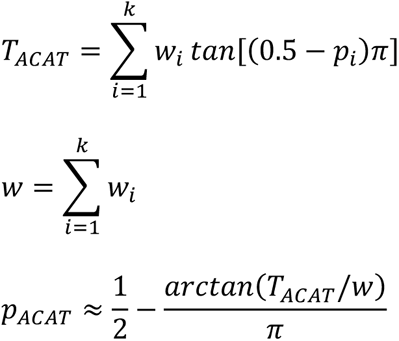

For each psychiatric disorder, we used the GWAS p-values for all MBSVs and weighed them by the absolute difference score. To further dissect the role of MBSVs in psychiatric disorders, we repeated the ACAT p-value aggregation on subsets of variants corresponding to each miRNA family the variants affected. In each case, four tests were conducted per GWAS/meta-analysis: all MBSVs, GTEx eQTL MBSVs, GTEx brain eQTL MBSVs, and PSYCHENCODE eQTL MBSVs. As such, a genome-wide significance threshold of 1.25x10^-8^ and a suggestive significance threshold of 2.5x10^-6^ were chosen as significance thresholds, based on typical GWAS significance levels (5x10^-8^ / 4 and 1x10^-5^ / 4 respectively).

### Gene- and gene set-level analysis of MBSV-affected pathways associated with psychiatric disorders

To investigate whether MBSVs were significantly aggregated within gene ontology gene sets associated with each disorder, we used MAGMA (v1.07) (de Leeuw et al., 2015). MBSVs were used to filter the GWAS variants supplied to MAGMA. Gene-level analyses and competitive gene set association tests were performed using gene ontology sets. Genes with a Bonferroni-corrected p-value < 0.05 were considered significant. Due to gene ontology sets not being strictly independent of one another, a relaxed significance threshold of an FDR-corrected p-value < 0.05 was applied to the gene set-level analyses. Gene set association was tested for all MBSVs, as well as MBSVs annotated as either GTEx eQTLs, GTEx brain eQTLs or PSYCHENCODE eQTLs.

### Meta-analysis of MBSV and MBSV target pathway associations across psychiatric disorders

To investigate whether there were general associations between MBSV-affected miRNA families and gene pathways across all nine disorders, we applied a meta-analysis technique. For the gene-set association meta-analysis, we used the --meta option in MAGMA. This option utilises the Stouffer’s weighted Z-score meta-analysis method. Briefly, p-values are converted to a standardised Z-score *via* the probit function. The Z-scores are then subsequently combined:

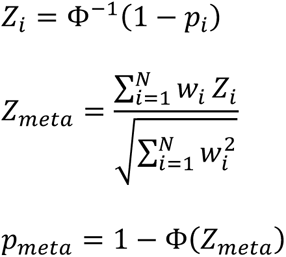

The Z-scores weights *w*_*i*_ are the sample sizes of the respective GWAS. We further employed this same method to meta-analyse the ACAT-generated p-values for the miRNA family associations.

### Analysis of difference score associations with GWAS effect size

To determine whether any specific miRNA families demonstrated a significant relationship between the difference score of each MBSV and the associated GWAS effect size for each disorder, we conducted a series of linear regression analyses. Effect sizes were standardised to be relative to the alternate allele by inverting the sign of the log-transformed values if the original GWAS data was calculated relative to the reference allele. We additionally investigated the relationship between the sign of the difference scores and effect sizes. For each disorder, we only investigated those miRNA families which showed a suggestive-significant ACAT-aggregated p-value (p < 2.5x10^-6^). Significance was determined by a Bonferroni-corrected p-value < 0.05.

### Comparison against non-psychiatric traits

To determine whether our findings were specific to psychiatric disorders or a more general phenomenon of human traits, we repeated the above analyses for several non-psychiatric traits, including body mass index (BMI) (Pulit et al., 2019), type 2 diabetes (both adjusted and unadjusted for BMI) (Mahajan et al., 2018), height (Yengo et al., 2018), and cardiovascular disease (CAD) (EPIC-CVD Consortium et al., 2017). To identify MBSVs for these traits, we used blood gene and miRNA expression as a reference; we further repeated the type 2 diabetes analyses using pancreas gene and miRNA expression as a reference. Similarly, in eQTL analyses, MBSVs were filtered for those annotated in the respective tissues in the GTEx database. Blood gene and miRNA expression data were acquired from RNA sequencing of peripheral blood mononuclear cells obtained from 15 healthy individuals in the Australian Schizophrenia Research Bank (ASRB) as detailed in a previous publication (M. P. Geaghan et al., 2019). Both gene and miRNA expression were filtered for a minimum CPM value of 0.381 and 9.21, respectively, in at least 75% of the samples, equating to a minimum count of 10 reads in the smallest libraries. Pancreas tissue expression data was obtained from publicly-released sequencing data of 5 healthy individuals as part of a study on pancreatic cancer (Müller et al., 2015). These data were available as averaged normalised expression values (NEV), and were filtered for an NEV greater than 10.

### Code availability

All R and bash scripts used for these analyses, as well as software version information is available at the following GitHub repository: https://github.com/mgeaghan/mbsv.

## Results

### MBSVs are enriched for higher functional scores

Utilising the dbMTS database, we investigated the union of all variants tested in each of the nine psychiatric disorder GWAS. From the list of allele-miRNA-mRNA interactions obtained from the database, we identified those interactions predicted by TargetScan and at least one of the other two algorithms (RNAhybrid and miRanda). We calculated scores representing the difference between the best reference allele interactions and the best alternate allele interactions, and discounted variants with small scores (absolute difference score < 0.2). In total, we identified a total of 8,334 variants, which we considered to be ‘true’ miRNA binding site variants (MBSVs). On average, MBSVs accounted for ∼0.067% of the SNVs in each GWAS (SD = 0.0025%), with between 3,123 and 3,995 independent (pruned) MBSVs out of 391,333 - 835,573 total variants (mean proportion MBSVs = 0.59%, SD = 0.12%) (Table 1). Similar proportions of MBSVs were found among the non-psychiatric traits (Supplementary Tables 52a-b).

We next aimed to assess whether MBSVs were enriched for functional annotation with the CADD (combined annotation dependent depletion) score. For every variant, we retrieved its CADD score, then compared the empirical cumulative distribution function of all variant CADD score ranks to the score ranks of MBSVs, GTEx eQTL MBSVs, GTEx brain eQTL MBSVs and PSYCHENCODE eQTL MBSVs. We obtained scores from both the most recent version of CADD (v1.6) as well as the original published version (v1.0). The use of both versions of the database was due to the addition of miRNA binding site scores from both TargetScan and mirSVR to the CADD v1.1 algorithm; as such, the later versions of the scores may be biased for MBSVs. For both the original and the latest CADD database versions, we observed a significant enrichment of CADD score ranks among all MBSV categories compared to all variants, except for PSYCHENCODE MBSVs, which were only nominally significant when using CADD v1.6 scores (Supplementary Table 1a, Supplementary Figure 1). To investigate whether this enrichment was specific to MBSVs or a general property of 3’ UTR variants, we also compared the ECDFs of MBSVs and eQTL MBSVs to non-MBSV 3’ UTR variants. Interestingly, there was a markedly reduced difference between MBSVs and other 3’ UTR variants (Supplementary Table 1b, Supplementary Figure 2). For CADD v1.6, the CADD score rank distributions of all MBSVs and all eQTL subsets of MBSVs were significantly different to that of non-MBSV 3’ UTR variants, with marginally elevated median score rank except for PSYCHENCODE eQTL MBSVs, which had a smaller median score rank. For CADD v1.0, all eQTL MBSV subsets were significantly different with smaller median score ranks compared to non-MBSV 3’ UTR variants.

We also found similar patterns of CADD score enrichment in blood and pancreas MBSVs (Supplementary Tables 53a-d, Supplementary Figures 9-12). Blood MBSVs (all MBSVs as well as eQTL and blood-eQTL MBSVs) displayed significantly higher median CADD score ranks (Supplementary Table 53a). The same result was seen in pancreas MBSVs (Supplementary Table 53c). Compared to 3’ UTR non-MBSVs (Supplementary Table 53b), all MBSVs displayed elevated v1.6 median CADD scores, eQTL MBSVs displayed elevated v1.6 median CADD scores but reduced median v1.0 CADD scores, and blood eQTL MBSVs showed reduced CADD scores from both versions of the database. For pancreas MBSVs compared to 3’ UTR non-MBSVs, v1.0 median CADD scores were significantly reduced.

### MBSVs are enriched in several psychiatric disorders

To determine if there was any association between MBSVs and each psychiatric disorder, we first analysed the empirical cumulative distribution functions of p-values for all independent variants in each GWAS dataset and compared these to the ECDFs of independent GWAS MBSVs. This revealed a significant enrichment of association between MBSVs and SCZ, BIP, MDD, ASD and AN after correction for multiple tests, with the median p-value for MBSVs being 0.063, 0.045, 0.037, 0.010 and 0.015 lower than the median p-value for all GWAS variants, respectively (Supplementary Table 2a, Supplementary Figure 3, Supplementary Figure 4). In support of these findings, we found that for SCZ there was a significant enrichment of genome-wide associated (p < 5x10^-8^) MBSVs (Supplementary Table 4a, Supplementary Figure 6), with 9 out of 3,502 (0.26%) MBSVs being genome-wide significant compared to 252 out of 594,558 (0.042%) of all independent non-MBSV GWAS variants (OR = 6.3; p = 2.3x10^-5^; FDR = 0.00028). No other GWAS displayed genome-wide significant MBSVs, although MDD displayed a nominal significance, with 3 out of 3,746 (0.080%)) MBSVs being genome-wide significant compared to 91 out of 707,808 (0.013%) (OR = 6.4 p = 0.013; FDR = 0.066). However, when a less rigorous variant-level p-value threshold was used (p < 1x10^-5^) as a measure of suggestive significance, BIP displayed significant enrichments of significant MBSVs (9/3995 (0.23%) vs 130/835573 (0.016%); OR = 15.5; p = 1.73x10^-8^; FDR = 4.14x10^-7^), in addition to SCZ (27/3502 (0.77%) vs 848/594558 (0.14%); OR = 5.6; p = 4.46x10^-12^; FDR = 1.95x10^-10^) and MDD (7/3746 (0.19%) vs 394/707808 (0.00058%); OR = 3.4; p = 0.0055; FDR = 0.033).

We next inquired as to whether MBSVs already annotated as eQTLs, and in particular as brain eQTLs were further enriched for association with these disorders. When considering MBSVs annotated as an eQTL for any tissue in GTEx, SCZ, BIP, MDD, ASD, AN, OCD, and TS all showed a significant enrichment of association (FDR < 0.05), with a greater difference in median p-value compared to all MBSVs (Supplementary Table 2a, Supplementary Figure 3, Supplementary Figure 4). SCZ, BIP, MDD and AN continued to show a significant enrichment when selecting only GTEx eQTLs for brain tissues, again with a greater difference in median p-value compared to all MBSVs and all GTEx eQTLs. SCZ, BIP and MDD were also significantly enriched for PSYCHENCODE eQTLs. SCZ and MDD retained a significant enrichment of genome-wide significant GTEx eQTLs (SCZ: 5/1023 (0.49%), OR = 11.7, p = 8.9x10^-5^, FDR = 0.00092; MDD: 3/1035 (0.29%), OR = 23.2, p = 0.00035, FDR = 0.0028); SCZ remained significant for GTEx brain eQTLs (SCZ: 2/212 (0.94%), OR=22.4, p = 0.0038, FDR = 0.025) (Supplementary Table 4a, Supplementary Figure 6).

Similar results were obtained when we compared the p-value distributions and proportions of significant variants between MBSVs and non-MBSV 3’ UTR variants (Supplementary Table 2b, 5b, Figure 1, Supplementary Figure 5). AN, BIP, MDD and SCZ displayed significant enrichments of association among all MBSVs; all but PTSD and ADHD further showed significant enrichment among all MBSV categories. Genome-wide significant MBSVs were significantly enriched in SCZ, and suggestive-significant MBSVs were enriched in SCZ and BIP (Figure 2).

**Figure 1.**
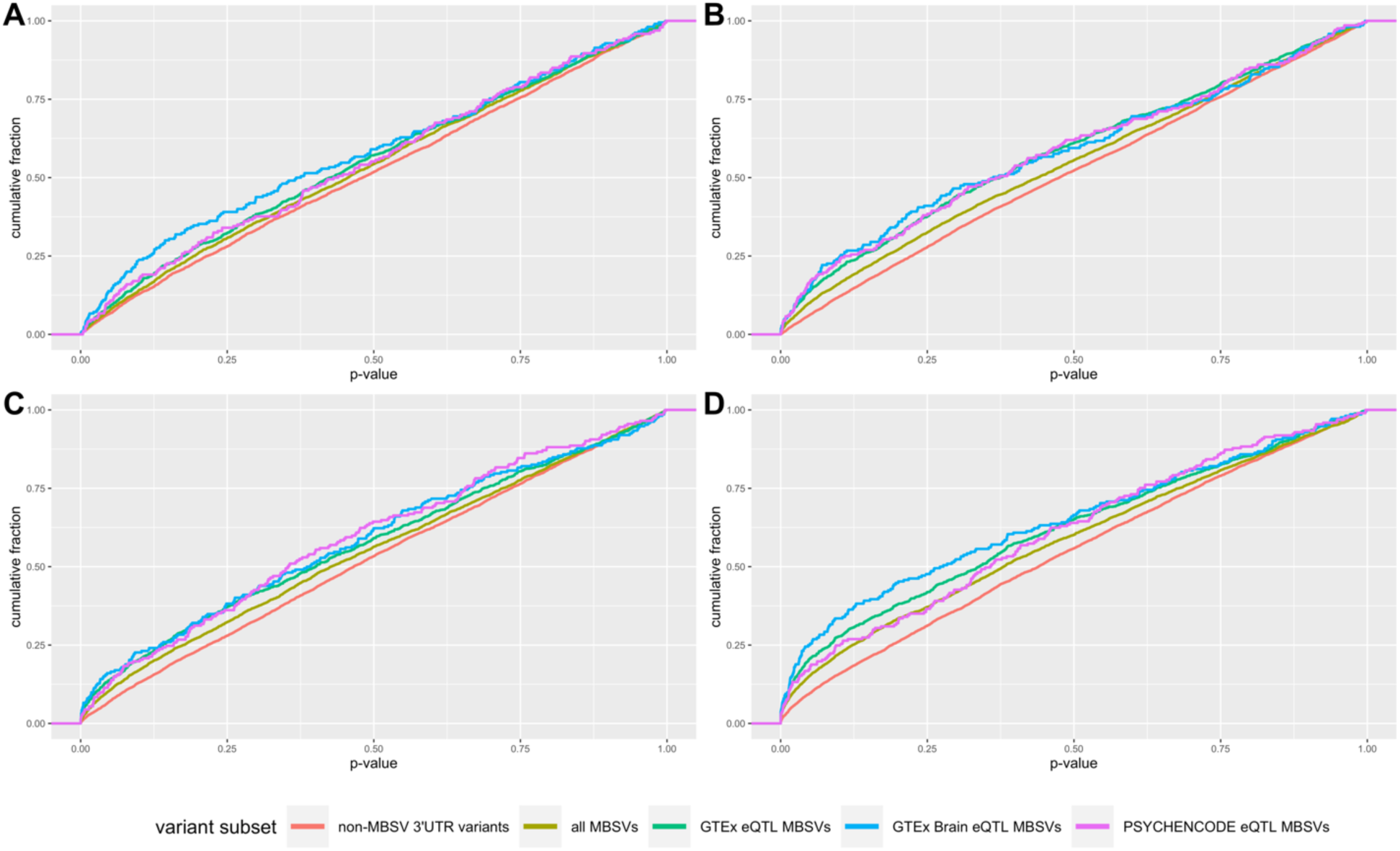
Distribution of p-values for AN, BIP, MDD, and SCZ. For each disorder, p-values for MBSVs and eQTL MBSVs were retrieved and compared to 3’ UTR-localised non-MBSVs with the Kolmogorov-Smirnov test. Median p-values were also compared. Significant (Benjamini-Hochberg FDR < 0.05) differences in p-value distributions of all MBSVs were identified in AN **(a)**, BIP **(b)**, MDD **(c)**, and SCZ **(d)**, with each of these disorders displaying a significant difference in the distribution of at least one eQTL MBSV category (GTEx eQTL MBSVs, GTEx brain eQTL MBSVs, or PSYCHENCODE eQTL MBSVs). Neither ADHD nor PTSD showed any significant differences, while ASD, OCD and TS showed a slight difference for GTEx eQTL MBSVs only (see Supplementary Figure 5). In each case of altered p-value distribution, the median p-value was lower for MBSVs compared to 3’ UTR-localised non-MBSVs.

**Figure 2.**
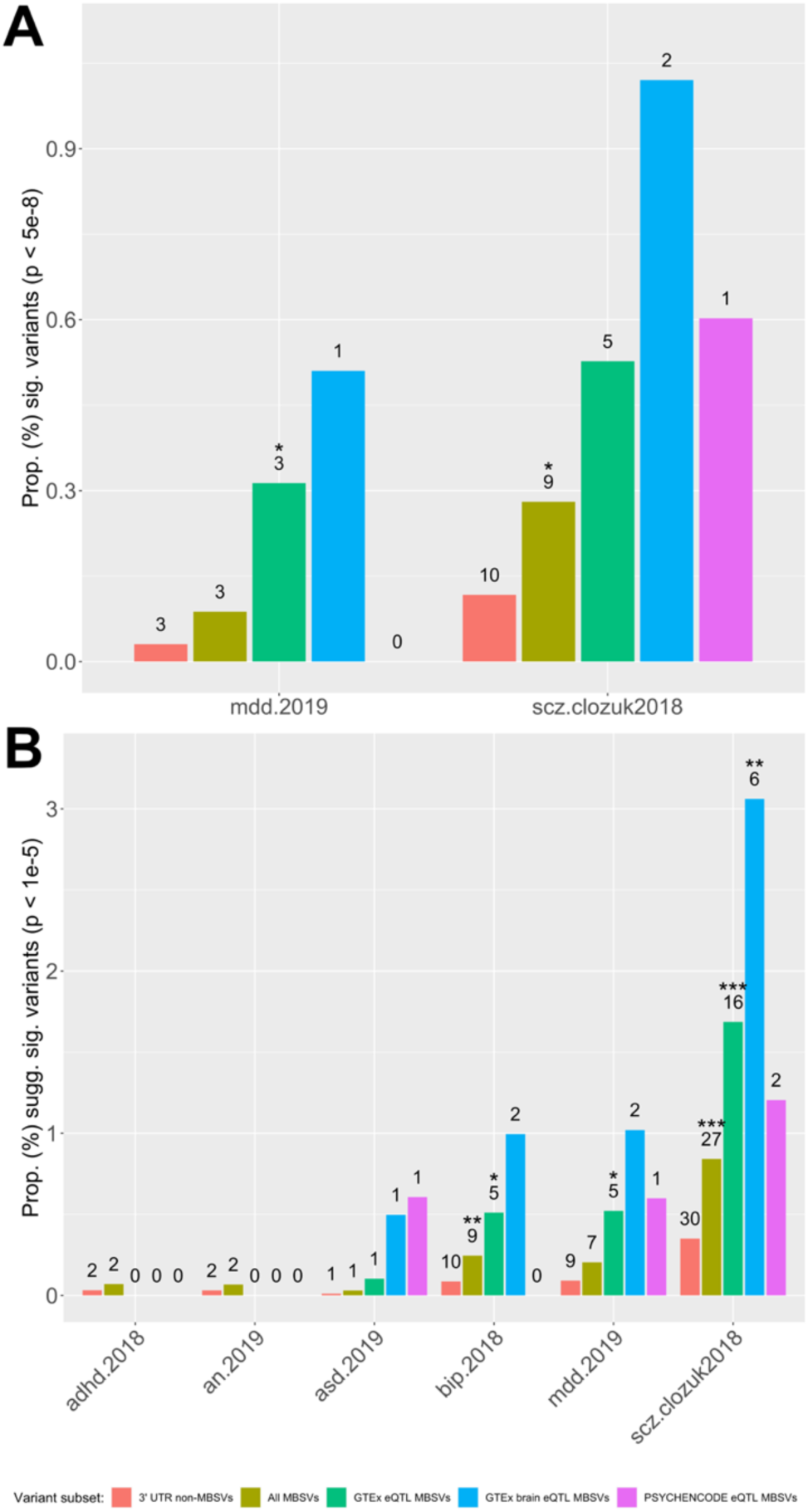
Proportions of strongly-associated variants in each disorder. Variants were filtered for genome-wide significance (p < 5x10^-8^) **(a)** or suggestive significance (p < 1x10^-5^) **(b)**. Proportions of all strongly-associated MBSVs (gold) and eQTL-annotated MBSVs (GTEx in green; GTEx brain in blue; PSYCHENCODE in pink) were compared to the proportion of strongly-associated 3’UTR-localised non-MBSVs (red) using Fisher’s exact test. Only disorders with at least one significant MBSV are shown. Numbers represent counts of significant variants. * = p < 0.05; ** = p < 0.01; *** = p < 0.001.

Additionally, we aggregated all MBSV p-values for each disorder using the ACAT method. Again, there was a significant association between MBSVs and SCZ and MDD (ACAT p < 1.25x10^-8^) (Supplementary Table 5, Figure 3). BIP and ADHD only showed nominal associations (p < 1x10^-3^). Furthermore, MBSV that were GTEx eQTLs were significantly associated with both SCZ and MDD, and GTEx brain eQTL MBSVs were associated with SCZ. Due to the significant genetic correlation between SCZ, MDD, BIP (SCZ-BIP: 68%; SCZ-MDD: 34%; MDD-BIP: 35%) (Brainstorm Consortium et al., 2018), and given that MBSVs were nominally associated with BIP, we also performed several meta-analyses of these p-values using Stouffer’s weighted Z-scoreethod: SCZ and MDD; SCZ and BIP; SCZ, MDD and BIP; and a final meta-analysis across all disorders. Each of these meta-analyses revealed a highly significant association (p < 1.25x10^-8^) of MBSVs, as well as GTEx eQTL and GTEx brain eQTL MBSVs (Supplementary Table 15).

**Figure 3.**
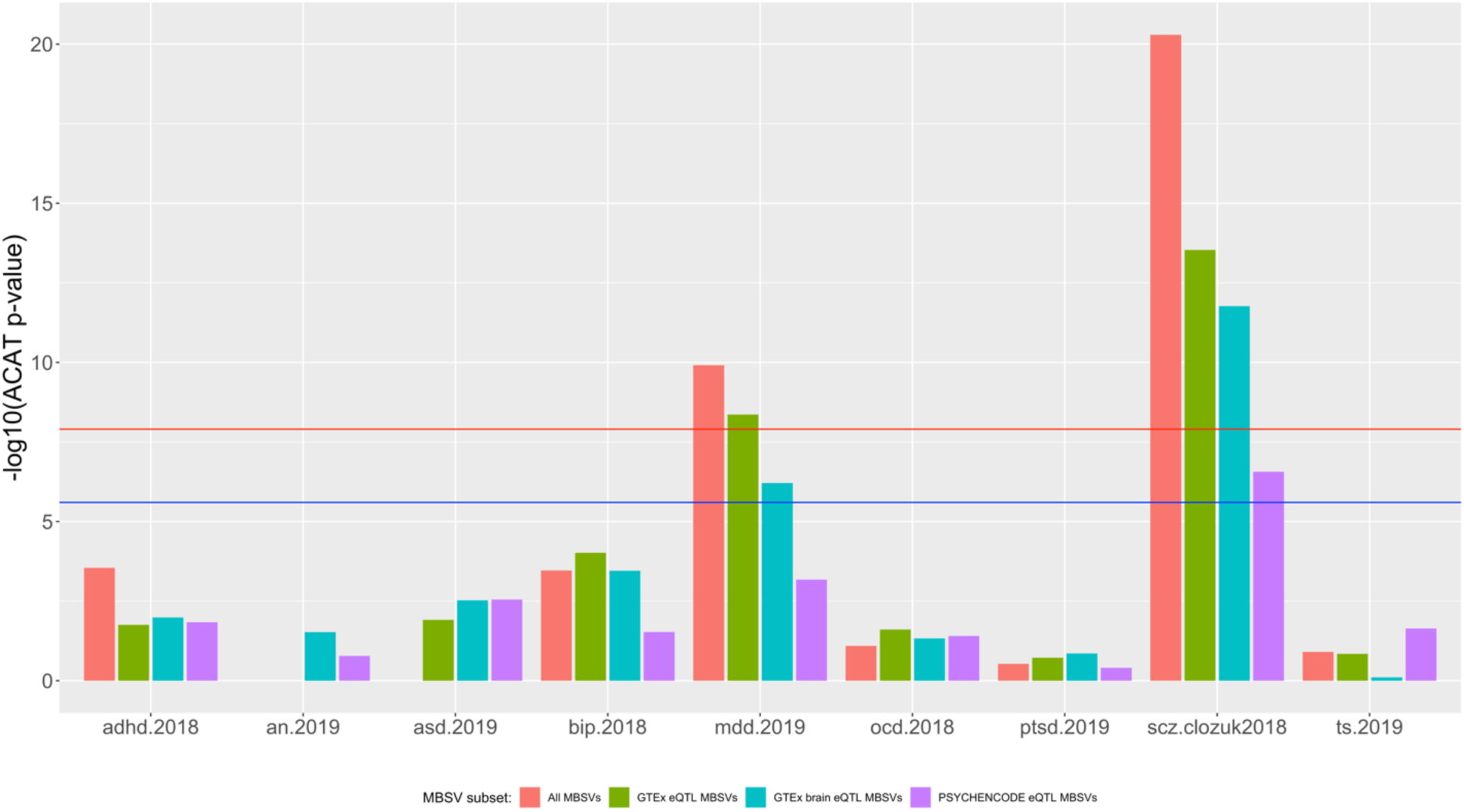
Aggregated p-values for MBSVs in each disorder. P-values for all MBSVs (red) and eQTL-annotated MBSVs (GTEx in green; GTEx brain in blue; PSYCHENCODE in pink) were aggregated using the ACAT method into one test statistic per disorder. Red line = genome-wide significance corrected for multiple tests (1.25x10^-8^); blue = liberal significance threshold corrected for multiple tests (2.5x10^-6^).

We also sought to determine if MBSVs displayed a significantly different effect size compared to all GWAS variants. We calculated the ECDF of the absolute log-transformed effect sizes (log-OR or beta) for MBSVs and all GWAS variants for each disorder. Interestingly, for each disorder, there was a significantly lower median absolute log effect size for MBSVs (p < 2.2x10^-16^), and this remained true for GTEx eQTLs, GTEx brain eQTLs and PSYCHENCODE eQTLs, suggesting that MBSVs had a smaller effect size than the average variant (Supplementary Table 3a, Supplementary Figure 7). These results were recapitulated when comparing MBSVs to non-MBSV 3’ UTR variants (Supplementary Table 3b, Supplementary Figure 8).

### MBSVs are also enriched in non-psychiatric traits

We found similar results when investigating the proportions of MBSVs in non-psychiatric traits, as well as the distributions of their p-values and absolute log-transformed effect sizes. Except for CAD, each trait was enriched for genome-wide- and suggestive-significant MBSVs and eQTL MBSVs compared to all non-MBSVs (Supplementary Table 56a, Supplementary Figure 17); height and T2D (both adjusted and unadjusted for BMI) were enriched for blood eQTL MBSVs, and both T2D GWAS were enriched for pancreas eQTL MBSVs (Supplementary Table 56c, Supplementary Figure 19). Genome-wide significant MBSVs were also enriched relative to 3’UTR non-MBSVs in both T2D GWAS and height (Supplementary Table 56b, Supplementary Figure 18), and eQTL MBSVs were enriched in both T2D GWAS. MBSVs below a p-value threshold of 1x10^-5^ were further enriched in both T2D GWAS, BMI, and height, with eQTL and blood eQTL MBSVs enriched in the two T2D GWAS.

For all MBSV subsets from both blood and pancreas, and for both comparisons against all non-MBSVs and all 3’UTR non-MBSVs, all tested non-psychiatric traits showed a significant difference in p-value distributions, with median p-values being lower than non-MBSVs in each case (Supplementary Tables 54a-d, Supplementary Figures 13-16). All comparisons of T2D MBSVs further showed a significant change and drop in median absolute log-effect sizes (Supplementary Tables 55a-d, Supplementary Figures 23-24). BMI MBSVs displayed marginally elevated absolute log-effect sizes relative to all non-MBSVs (Supplementary Table 55a). CAD MBSV effect sizes were significantly smaller in all comparisons, while height showed a small elevation in the effect sizes of all MBSVs and eQTL MBSVs when compared against all non-MBSVs.

When aggregating p-values of MBSVs for each trait, we found a significant association between all MBSVs and all traits except CAD; similarly, we found a significant association between all eQTL MBSVs and all traits except CAD, and a significant association between blood eQTL MBSVs and BMI and height (Supplementary Table 57a, Supplementary Figure 21). We also found a significant association between pancreas eQTL MBSVs and T2D (BMI adj.) (Supplementary Table 57b, Supplementary Figure 22).

### Variants affecting binding of specific miRNA families are associated with psychiatric disease

We next aggregated MBSV p-values separately for individual miRNA families in each disorder. We identified several miRNA families reaching genome-wide significance (p < 1.25x10^-8^) in MDD and SCZ (Supplementary Table 10, Supplementary Table 13). Four miRNA families (miR-335-5p, miR-21-5p/590-5p, miR-361-5p and miR-577) were highly significantly associated (p < 1.25x10^-8^) with both disorders, with a further 3 families showing association with MDD (miR-31-5p, miR-151-3p, and miR-654-3p) and another 14 with SCZ (miR-22-3p, miR-23-3p, miR-450b-5p, miR-193a-5p, miR-770-5p, miR-338-3p, miR-409-3p, miR-323b-3p, miR-194-5p, miR-28-3p, miR-500a-3p, and miR-129-5p). At a less rigorous level of significance (p < 2.5x10^-6^), 3 more miRNAs were significant in MDD, and 10 more were significant in SCZ, including the previously SCZ-associated miR-132-3p/212-3p family. In total, 26 miRNAs passed a significance threshold of 2.5x10^-6^ in SCZ, and 10 passed this threshold for MDD. We further meta-analysed the results for SCZ-MDD, SCZ-BIP, SCZ-MDD-BIP, and all disorders. At a significance threshold of 2.5x10^-6^, 24 miRNA families were significant in all meta-analyses (Supplementary Table 16-19). The families miR-335-5p, miR-21-5p/590-5p, miR-361-5p and miR-577 were consistently genome-wide significant (p < 1.25x10^-8^) and amongst the top 9 families in each analysis. Similarly, 16 families were consistently significant when considering GTEx eQTL MBSVs, with miR-21-5p/590-5p, miR-361-5p and miR-577 consistently the top 8 families. Few families remained significant when considering brain or PSYCHENCODE eQTL MBSVs.

### Non-psychiatric traits display associations with miRNA family-specific MBSVs

Aggregation of MBSV p-values into individual miRNA families revealed that, except for CAD, multiple miRNA families were associated with each non-psychiatric trait (Supplementary Tables 58-64); for CAD only miR-22-3p eQTL and blood eQTL MBSVs were nominally significantly associated (Supplementary Table 61). For T2D (BMI unadj., blood MBSVs), significant associations were found with miR-485-5p, miR-181-5p, miR-139-5p, and miR-143-3p (Supplementary Table 58). T2D (BMI adj., blood MBSVs) was associated with miR-485-5p (Supplementary Table 59). T2D (BMI unadj., pancreas MBSVs) was associated with miR-129-5p, miR-455-5p, miR-181-5p, miR-660-5p, miR-139-5p, miR-200bc-3p/429, miR-217, miR-1185-5p, miR-143-3p, and miR-199-5p (Supplementary Table 63). T2D (BMI adj. pancreas MBSVs) was associated with miR-129-5p and miR-455-5p (Supplementary Table 64). Interestingly, height was associated with the greatest number of miRNA families with 91, 67 and 21 among all blood MBSVs, GTEx eQTL MBSVs, and blood eQTL MBSVs respectively (Supplementary Table 62). Among the most significant of these miRNA families were miR-144-5p, miR-576-5p, miR-103-3p/107, and miR-423-5p.

### Relationship between MBSV difference scores and GWAS effect sizes

We next investigated whether the difference scores for MBSVs in each associated miRNA family in each psychiatric disorder demonstrated a relationship with the variants’ effect sizes. We only considered miRNA families passing an ACAT p-value threshold of 2.5x10^-6^. The effect sizes were standardised such that they were all relative to the alternate allele, as the difference scores were also calculated relative to the alternate allele. A linear regression analysis revealed no significant relationship between difference scores and effect sizes for any miRNA family in any disorder. However, we also investigated whether the sign of the difference scores were associated with effect size. In this analysis, we identified one miRNA family – miR-323b-3p – in SCZ that demonstrated a significant negative relationship – i.e. a positive difference score (indicating a loss of binding affinity in the presence of the alternate allele) was associated with a more negative effect size, and vice versa (p = 0.0015, Bonf. p = 0.045) (Figure 4, Supplementary Tables 46-51).

**Figure 4.**
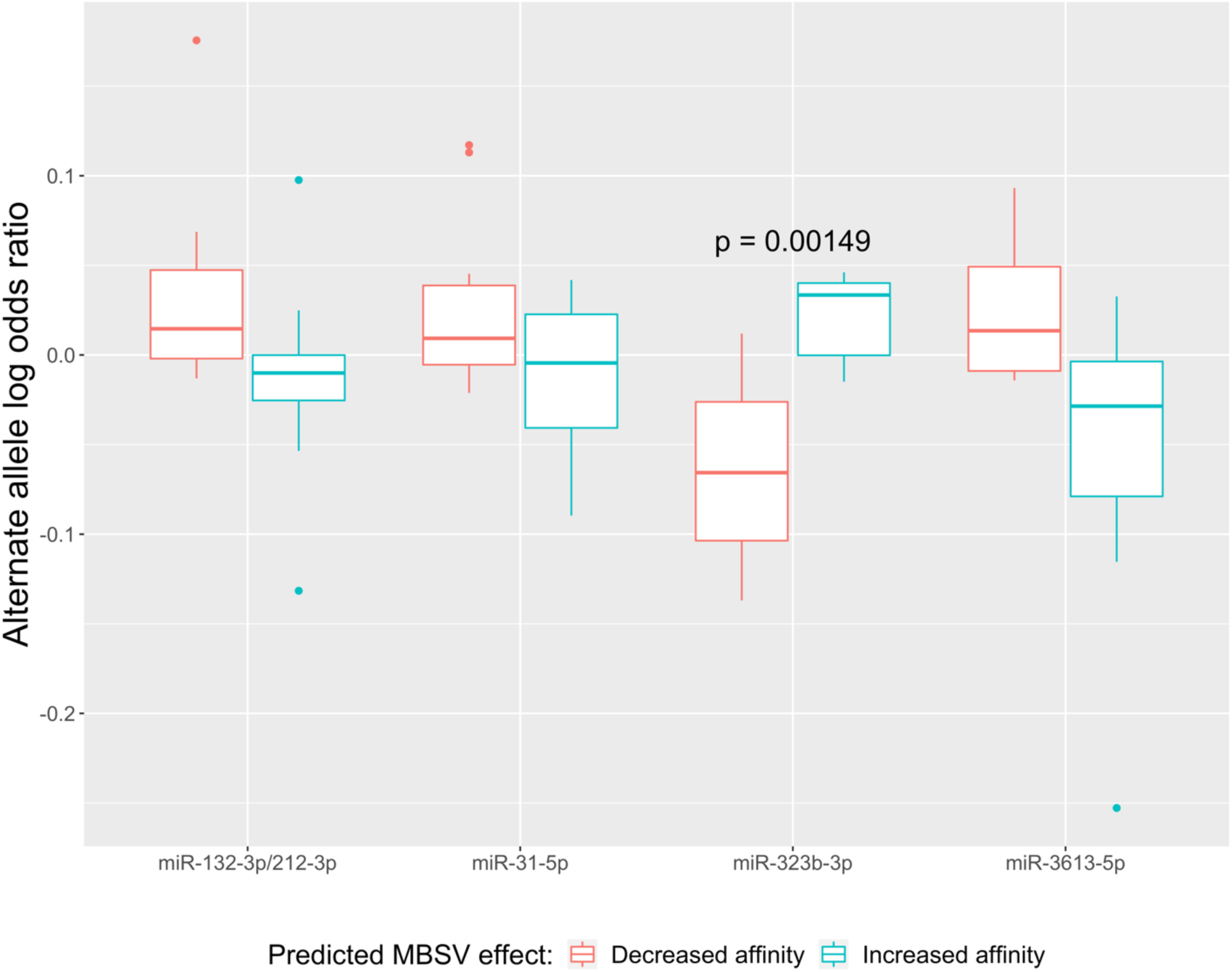
Relationship between the sign of MBSV family difference scores and SCZ effect sizes. Effect sizes were the log-transformed odds ratios from the 2018 SCZ GWAS (Pardiñas et al., 2018), and were standardised to be relative to the alternate allele, as all difference scores were calculated for the alternate alleles. A linear regression model was fitted to determine whether difference score sign – positive scores representing lower binding affinity and vice versa – was significantly associated with effect size. Shown are the results for the four nominally significant (p < 0.05) miRNA families. A significant relationship (Bonferroni-corrected p-value < 0.05) was found only for miR-323b-3p in SCZ.

We additionally found a few significant relationships between difference scores and effect sizes of non-psychiatric MBSVs (Figure 5). Height demonstrated a significant negative relationship between the difference score for miR-1343-3p and the alternate allele effect size, i.e. loss of binding affinity in the presence of the alternate allele was associated with a more negative effect size (p = 0.000171) (Supplementary Table 83). T2D (BMI adj., blood MBSVs) showed a negative relationship for miR-139-5p between the sign of the difference score and alternate allele effect size (p = 0.00149) (Supplementary Table 87), as did BMI for miR-142-5p (p = 0.000776).

**Figure 5.**
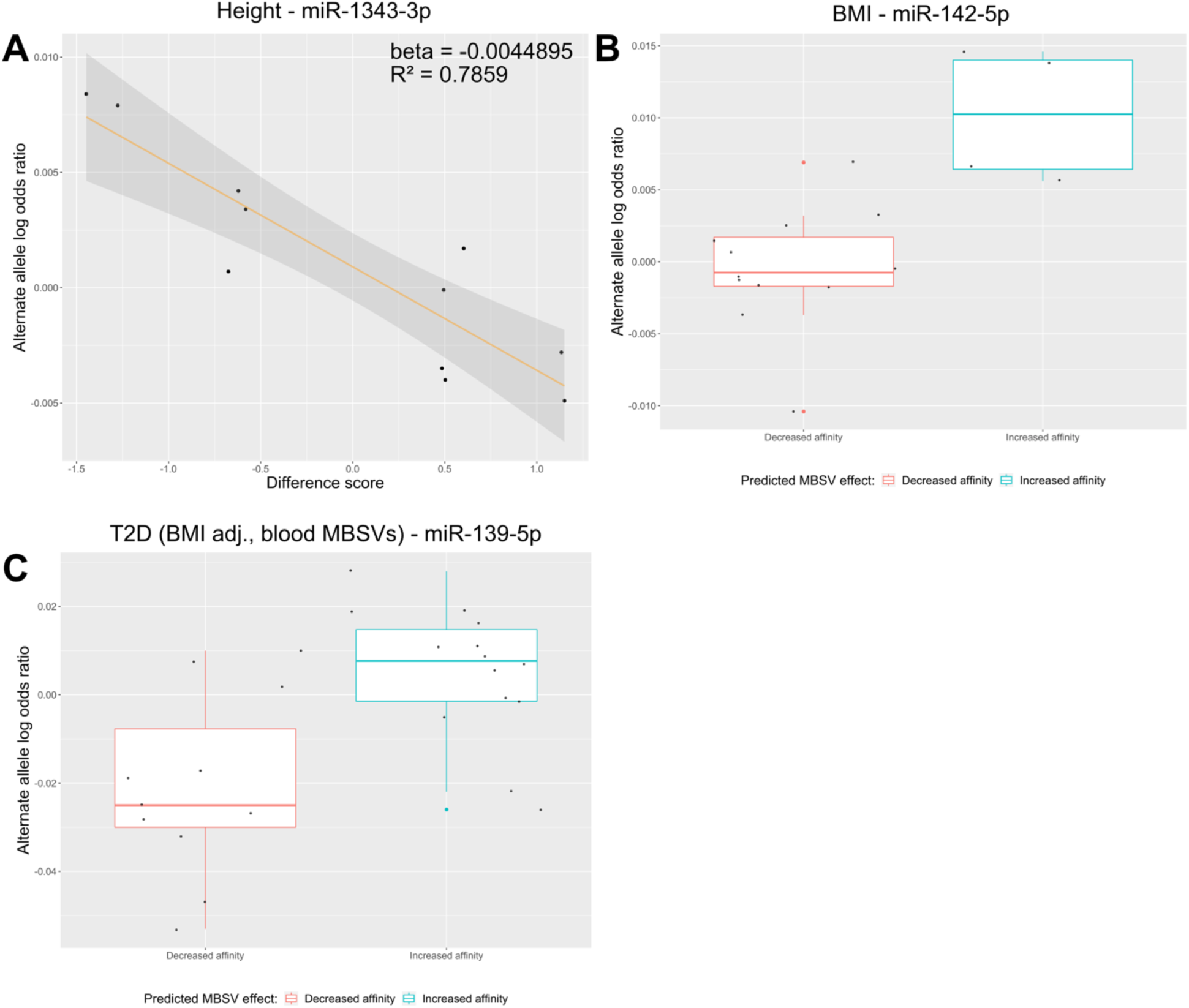
Relationship between the MBSV family difference scores and effect sizes in HEIGHT **(a)**, BMI **(b)**, and T2D (BMI adj.) **(c)**. Effect sizes were the log-transformed odds ratios from each GWAS, and were standardised to be relative to the alternate allele, as all difference scores were calculated for the alternate alleles. A linear regression model was fitted to determine whether either difference score or the sign of the difference score – positive scores representing lower binding affinity and vice versa – was significantly associated with effect size. For HEIGHT **(a)** there was a significant negative linear relationship between the difference scores for miR-1343-3p family MBSVs and effect size. For BMI **(b)** and T2D (BMI adj.; blood MBSVs) **(c)** there was a significant difference in effect size between MBSVs predicted to increase binding affinity and those predicted to decrease binding affinity of miR-142-5p and miR-139-5p, respectively, with increased affinity in the presence of the alternate allele associated with an elevated log-odds ratio. These results remained significant after Bonferroni correction for multiple tests.

### Analysis of gene sets affected by MBSVs

To identify whether MBSVs had any aggregated effects within disease-relevant pathways, we utilised MAGMA to perform gene-set association analyses for each GWAS, subset by MBSVs. Gene sets were all gene ontology sets obtained from the molecular signatures database (MSigDB). At an FDR threshold of 0.05, we identified 2, 3, and 1 gene sets in AN, SCZ, and TS, respectively, that were significantly enriched for association with MBSVs. We further identified 6 and 12 gene sets in ASD and TS significantly enriched for association with PSYCHENCODE eQTL MBSVS, and an additional 2 and 1 gene sets in SCZ significantly enriched for eQTL and brain eQTL MBSV association, respectively (Supplementary Tables 20-28). Of particular interest, among all MBSVs in SCZ, there was a significant association within the gene set “*Regulation of long term synaptic depression*” (p = 0.000018, FDR = 0.043); among eQTL MBSVs in SCZ, “*Negative regulation of synapse organization*” was significantly associated (p = 0.000013, FDR = 0.037). In TS, two nervous system-relevant gene sets – “*Neural crest cell migration*” (p = 2.31x10^-5^, FDR = 0.028) and “*Neural crest cell differentiation*” (p = 2.31x10^-5^, FDR = 0.028) – were also significantly associated among PSYCHENCODE eQTL MBSVs. At the gene level, several genes affected by MBSVs were significantly associated with ADHD, AN, ASD, BIP, MDD, and SCZ; eQTL, brain eQTL, and PSYCHENCODE eQTL MBSVs also were significantly associated with ASD, BIP, MDD and SCZ (FDR < 0.05) (Supplementary Tables 33-41). Interestingly, an MBSV eQTL in the miRNA biogenesis gene *AGO1* was significantly associated with SCZ (p = 7.9x10^-6^, Bonf. p = 0.038).

Again, we investigated whether any gene ontology sets displayed joint association when the gene-level results from MAGMA for these disorders were meta-analysed. We conducted four different meta-analyses: SCZ and BIP; SCZ and MDD; SCZ, MDD and BIP; and all disorders (Supplementary Tables 29-32). Among several findings in the SCZ-BIP analysis, “*Protein localization to synapse*” (p = 1.35x10^-5^, FDR = 0.026) and “*Negative regulation of synapse organisation*” (p = 8.15x10^-6^, FDR = 0.025) were significantly enriched for association with MBSVs and eQTL MBSVs, respectively. In the all-disorder analysis, “*Nerve growth factor binding*” was among several gene sets enriched for association with MBSVs (p = 1.51x10^-5^, FDR = 0.033).

For non-psychiatric traits, we found that each trait had at least one significantly associated gene set. For T2D (BMI unadj., blood MBSVs) “*Semi lunar valve development*” was significantly associated with eQTL MBSVs; for T2D (BMI adj., blood MBSVs), the same gene set, as well as “*Cardiac septum morphogenesis*” and *“Heart valve development*” were significantly associated with eQTL MBSVs. In the context of pancreas MBSVs, *“Cardiac atrium development*”, and *“Semi lunar valve development*” were associated with eQTL MBSVs in T2D (BMI unadj.) and “*Triglyceride lipase activity*” was further associated with all MBSVs; and for T2D (BMI adj., pancreas MBSVs), eQTL MBSVs were associated with “*Semi lunar valve development*”, “*Cardiac atrium development*”, “*Heart valve development*” and “*Cardiac chamber morphogenesis*”, and pancreas eQTL MBSVs were associated with “*Cardiac atrium development*”, “*Heart morphogenesis*” and “*Cardiac chamber morphogenesis*”. All MBSVs in BMI were significantly associated with “*Methylation dependent chromatin signalling*”, while blood eQTL MBSVs were associated with “*Lipid localization*”. In CAD, all MBSVs were associated with “*Positive regulation of dendrite development*”. Finally, in height, all MBSVs were associated with “*Response to vitamin A*”, “*Glucose metabolic process*”, and “*Dedifferentiation*”.

## Discussion

In this study, we aimed to investigate whether common genetic variation within miRNA binding sites contributes towards the pathophysiology of psychiatric disorders. Psychiatric disorders are typically complex, polygenic conditions with many small common variants, each with relatively small effect sizes, contributing towards overall risk. The majority of risk variants tend to lie outside coding regions and are thought to instead affect gene regulatory sequences, such as promoters and enhancer regions (Bray & O’Donovan, 2019). Concordant with this hypothesis, miRNA are increasingly being studied for their roles in psychiatric disease, as biomarkers, and as potential novel treatment targets (Natalie J. Beveridge & Cairns, 2012; M. Geaghan & Cairns, 2015; B. H. Miller & Wahlestedt, 2010). Both genetic and experimental biological studies have demonstrated a significant role for miRNAs in brain development, neuronal function and the pathophysiology of psychiatric disease. For example, the genetic locus containing the miR-137 host gene *MIR137HG* is host to one of the most significant common genetic associations with schizophrenia (Pardiñas et al., 2018; Schizophrenia Working Group of the Psychiatric Genomics Consortium, 2014). Further biological evidence suggests that this miRNA is responsible for regulating the formation and maturation of synapses, and its overexpression has been observed to result in reduced synapse formation and synaptic transmission (He et al., 2018). It has also been implicated in other psychiatric and neurological conditions, including autism spectrum disorders, Huntington’s disease and Rett syndrome (Mahmoudi & Cairns, 2017). Other miRNAs, as well as their biogenesis machinery have frequently been implicated in schizophrenia and other psychiatric disorders, as well as in the regulation synaptic and neuronal function, such as the miR-132/212 cluster and miR-185 (N. J. Beveridge et al., 2010; Earls et al., 2012; M. Geaghan & Cairns, 2015; M. P. Geaghan et al., 2019; B. H. Miller et al., 2012). We hypothesised that miRNA binding site sequences, present within 3’ UTRs of protein-coding genes, may represent yet another class of regulatory sequences that are affected by common variants that predispose towards psychiatric disease. While previous studies have investigated the role of these variants within schizophrenia (Devanna et al., 2018; Hauberg, Holm-Nielsen, et al., 2016; Hauberg, Roussos, et al., 2016) a large study of MBSVs across several psychiatric disorders has not been performed to date. To this end, we identified variants predicted to affect miRNA binding sites. As miRNA prediction algorithms suffer from relatively high false-positive rates (Lewis et al., 2005), we only considered binding sites predicted by TargetScan and at least one of the other two algorithms employed by dbMTS (RNAhybrid and miRanda). We constructed a difference score based on the TargetScan scores by subtracting the best (most negative) score for the reference allele from the best score for the alternate allele. This differed from the pre-calculated difference score supplied in the dbMTS database. We reasoned that the most conservative score would reflect the difference between the miRNA-mRNA interactions with the highest affinities for each allele, in contrast to the dbMTS approach of using the maximum absolute difference in scores. We further filtered out variants with small difference scores (absolute score < 0.2). The remaining 9,367 variants were considered ‘true’ MBSVs.

Five psychiatric disorders including SCZ, BIP, MDD, ASD and AN, were significantly enriched with independent MBSV association compared to all independent variants. In SCZ and MDD these findings were also supported by a significant enrichment of genome-wide associated MBSVs. This suggests that MBSVs are a significant source of genetic risk in both these disorders. We further dissected our data by investigating only MBSVs annotated as eQTLs and specifically brain eQTLs for the gene in which they are present. While there are significantly fewer MBSVs that are also eQTLs and brain eQTLs, we still observed a significant enrichment of association of these variants compared to all non-MBSVs for SCZ and MDD and found a significant enrichment of genome-wide associated MBSVs in SCZ. It should also be noted that the BIP, ASD and AN GWAS were supported by smaller sample sizes compared to MDD and SCZ (Grove et al., 2019; Howard et al., 2019; Pardiñas et al., 2018; Stahl et al., 2019; Watson et al., 2019), and as such, analysis of future GWAS may demonstrate a more significant enrichment of genome-wide associated MBSVs.

An interesting finding was that MBSVs typically display smaller effect sizes than other variants; this effect was further enhanced when considering only MBSVs annotated as eQTLs or brain eQTLs. This is not completely surprising, since miRNAs are typically subtle regulators of gene expression; their role is typically framed as a “fine-tuner” of protein expression within the cell (Selbach et al., 2008). Furthermore, a typical mRNA will possess several miRNA binding sequences within its 3’ UTR; thus, a single dysfunctional miRNA may be compensated for by other miRNAs, reducing the effect size of any one MBSV. Environmental factors, such as cellular and physical stress, can also influence miRNA expression (Lukiw & Pogue, 2007; Mannironi et al., 2013; Rocha, 2007), and thus there may be significant genotype-by-environment (GxE) interactions present, whereby MBSVs are only pathologically relevant within the context of other environmental risk factors for psychiatric disease. As a result, while common variation within these binding regions may contribute significantly towards disease risk, they may be less potent than other regulatory sequence variation. To add further support to this assertion, we observed that MBSVs tended to possess higher CADD score ranks when compared against all variants. The tendency towards higher CADD score ranks may support our hypothesis that these variants are functionally important. A significant caveat with the CADD scores was that they are calculated based on many functional annotations of known SNVs, and from v1.1 onwards these annotations include the TargetScan and mirSVR (Betel et al., 2010) target prediction algorithms. Thus, these scores are likely biased towards MBSVs. We therefore also utilised the original published CADD scores (v1.0) which did not include these annotations. This analysis returned similar results; however, even the original published version of CADD included 3’ UTR annotations, as this region of an mRNA is important for many forms of post-transcriptional regulation other than miRNA binding, such as RNA binding proteins (Berkovits & Mayr, 2015; Kuersten & Goodwin, 2003; Wilkie et al., 2003). As such, while the enrichment of CADD ranks may suggest a possible functional role of MBSVs, this result must be interpreted with caution.

By aggregating p-values assigned to each MBSV for each disorder, we identified a highly significant association between this class of variants and SCZ and MDD. Several specific miRNA families also showed association with both SCZ and MDD. Perhaps the most interesting finding was a strong association between MBSVs affecting the miR-132-3p/212-3p family and SCZ; the expression of this miRNA family has previously been associated with SCZ, and it is known to play a significant role in neuronal function. Specifically, miR-132-3p/212-3p targets are enriched for pathways associated with synaptic plasticity and neuronal migration pathways such as Reelin signalling (B. H. Miller et al., 2012). Furthermore, miR-132-3p expression increases during early post-natal development in mice and is positively regulated by NMDAR activation (B. H. Miller et al., 2012). In the current study, this miRNA family retained this suggestive association when meta-analysed across SCZ, MDD and BIP, as well as in the all-disorder meta-analysis. This result represents a significant finding that strengthens the support for a role of the miR-132/212 cluster in the pathophysiology of schizophrenia.

Interestingly, little evidence for association was observed for MBSVs affecting the miR-137 family in any disorder. This miRNA has been of particular interest, particularly within the context of SCZ, primarily due to the highly significant genetic association within its genetic locus (Pardiñas et al., 2018; Schizophrenia Working Group of the Psychiatric Genomics Consortium, 2014). This does not suggest that this miRNA has been erroneously associated with SCZ; rather, it only suggests that genetic variants affecting miR-137 binding do not significantly affect SCZ risk. As discussed above, various factors, including compensation by other miRNA binding sites, and genotype-by-environment interactions, may mitigate the effects of individual variants on the binding of any single miRNA. Instead, variants affecting miR-137 expression and the expression of its target genes may be more important for determining disease risk. Indeed, miR-137 targets (as predicted by TargetScan v5) were enriched for genome-wide significant associations in the 2014 PGC SCZ GWAS (Schizophrenia Working Group of the Psychiatric Genomics Consortium, 2014). Furthermore, a variable-number tandem repeat (VNTR) present in the *MIR137HG* locus has been demonstrated to regulate miR-137 expression, with shorter repeats linked to higher miR-137 expression and increased SCZ risk (Pacheco et al., 2019). However, it should also be emphasised that this locus also contains the *MIR2682* miRNA gene, the *MIR137HG* long non-coding RNA that hosts these two miRNAs, and *DPYD*. Interestingly, *DPYD* (dihydropyrimidine dehydrogenase) has previously been linked to autism (Ben-David et al., 2011; Carter et al., 2011). As such, it is also possible that miR-137 is not the major causal factor influencing risk for SCZ.

Interestingly, the miRNA family miR-335-5p was consistently the top-most associated family when aggregating MBSV p-values. This miRNA has previously been associated with depression, whereby it was found to be downregulated in peripheral blood mononuclear cells (PBMCs) in individuals with MDD, and was further found to regulate the depression-associated glutamate metabotropic receptor 4 (*GRM4*) and was upregulated in response to treatment with the antipsychotic citalopram (J. Li et al., 2015). This miRNA was also among those upregulated in a previous study from our laboratory (N. J. Beveridge et al., 2010). Thus, this miRNA may be particularly important for these conditions, and represents an interesting focus for future research into miRNAs in psychiatric disorders and their treatment.

We also found evidence for a relationship between MBSVs affecting miR-323b-3p binding and SCZ risk. The negative association between miR-323b-3p difference score and SCZ effect size suggests that increased binding affinity for this miRNA may be associated with elevated risk for SCZ. This miRNA has not been thoroughly studied in the context of psychiatric disease; the current literature does suggest it has a significant role in promoting neuronal apoptosis in the context of ischemia by targeting *BRI3* and *SMAD3* mRNA (Che et al., 2018; Yang et al., 2015). Similarly, it has been shown to inhibit proliferation and induce apoptosis in medulloblastoma, also *via* targeting *BRI3* (Zhang et al., 2017). As such, it is possible that this miRNA is involved in regulating the proliferation of cells in the developing brain. Importantly, mature miRNAs in this family are enriched in the rat brain (Kim et al., 2004), and are located within a maternally-imprinted cluster of miRNAs in the DLK1-DIO3 at 14q32 locus that were downregulated in SCZ and a two-hit rat model of maternal immune activation and adolescent cannabis exposure (Gardiner et al., 2012; Hollins et al., 2014). Furthermore, this miRNA family has been shown to regulate *FMR1* gene expression – a gene vital for dendritic spine and synapse development, and loss of function of which results in the debilitating neuropsychiatric disorder Fragile X syndrome (Nimchinsky et al., 2001; Yi et al., 2010). Together, this evidence suggests that this miRNA is important in neuronal development and function, and its role in SCZ is worthy of further study.

At the gene set level, we found multiple gene sets relating to neuronal function and nervous system development which were associated with SCZ and TS, respectively. Meta-analysis of SCZ and BIP further identified a significant association of “*Protein localization to synapse*” being observed among MBSVs. These results further support our hypothesis of the involvement of MBSVs in psychiatric disease and neuronal function and suggests that this class of variants may have a significant influence over the expression of genes involved in synaptic function. This fits with previous research suggesting that several psychiatric disease-associated miRNAs are potent regulators of synaptic activity and neuronal function. For example, the mature miRNAs arising from the miR-132/212 cluster have been demonstrated to influence basal synaptic transmission and aspects of long-term potentiation (LTP) in the hippocampus and neocortex in mice (Remenyi et al., 2013), and miR-132 specifically has been shown to regulate dendritic complexity and spine density, with subsequent effects on GABAergic and glutamatergic activity in olfactory bulb neurons (Pathania et al., 2012). Similarly, miR-137 has been shown to affect LTP, the synaptic vesicle pool, and aspects of learning and memory in mice (Siegert et al., 2015). It has also been shown that certain miRNAs, such as miR-134, can themselves be localised to the synapse (Schratt et al., 2006), and their biogenesis and silencing capacities can even be regulated by synaptic activity (Ashraf et al., 2006), thus allowing for fine control over protein synthesis at the synapse in response to neuronal stimuli. Our results add a new level of complexity to this regulatory system, suggesting that genetic variants affecting miRNA function may also influence processes of synaptic protein expression in psychiatric disorders.

We also repeated these analyses with non-psychiatric human traits, including type 2 diabetes, BMI, height, and cardiovascular disease. In most cases, we found a similar trend with a general enrichment of association between MBSVs and each trait and significant associations between specific families of MBSVs and each trait. Interestingly, we also observed significant associations between each trait and relevant MBSV-affected gene sets; for example, “*Lipid localization*” was significantly associated with blood eQTL MBSVs in BMI. These results suggest that the importance of MBSVs to the genetics of human traits is not specific to psychiatric disorders, but may be a more general phenomenon. This is not entirely unexpected given that miRNAs have been shown to be functionally significant in a plethora of human traits, conditions, and diseases, such as various types of cancer, neurological diseases such as Alzheimer’s and Huntington’s disease, and each of the non-psychiatric traits we have studied here (M. Geaghan & Cairns, 2015; Johnson et al., 2008; Lee et al., 2011; Swarbrick et al., 2019; Tong & Nemunaitis, 2008; W.-X. Wang et al., 2011). Thus, the methods employed in this study can be used to investigate the roles of MBSVs (and other non-coding regulatory regions) in a wide range of human traits, and could be useful in developing novel hypotheses and treatments for human diseases beyond psychiatric disorders.

Together, these results represent promising support for the role of MBSVs in regulating neuronal function in psychiatric disorders. However, the relatively small number of MBSVs compared to all common variants typically assessed in GWAS, and the difficulty of confidently predicting true miRNA binding sites means that the power to detect the effects of these variants is relatively low. Thus, repeating these analyses upon release of larger GWAS summary statistics, as well as in GWAS of different ethnic backgrounds, and upon the development of improved miRNA target prediction algorithms will be vital to the further exploration of the roles of MBSVs in complex disease. This approach can also be applied to other complex disorders for which GWAS summary data are available. While this study focussed on common variation, we suspect that rare MBSVs with greater effect sizes exist and that future studies focussing on these may shed more light on the influence of this class of regulatory variants in complex diseases. Nevertheless, these results suggest that variants that influence the binding of miRNAs to the 3’ UTRs of mRNAs are a significant source of genetic risk for complex disorders, and in the case of psychiatric disorders they may be affecting normal neuronal function by disrupting the localisation of proteins to the synapse. Further study of these variants may help to elucidate the complex molecular landscape of psychiatric and other complex disorders that ultimately may drive the development of novel treatments.

## Acknowledgements

This study was supported by a NARSAD Young Independent Investigator Grant (MC) and National Health and Medical Research Council (NHMRC) project grants (1067137, 1147644, 1051672). MG was supported by University of Newcastle RHD scholarship. MC is supported by an NHMRC Senior Research Fellowship (1121474) and a University of Newcastle Faculty of Health and Medicine Gladys M Brawn Senior Fellowship.

## Notes

### Competing Interest Statement

The authors have declared no competing interest.

