## Supplementary figures for "MicroRNA binding site variation is enriched in psychiatric disorders"

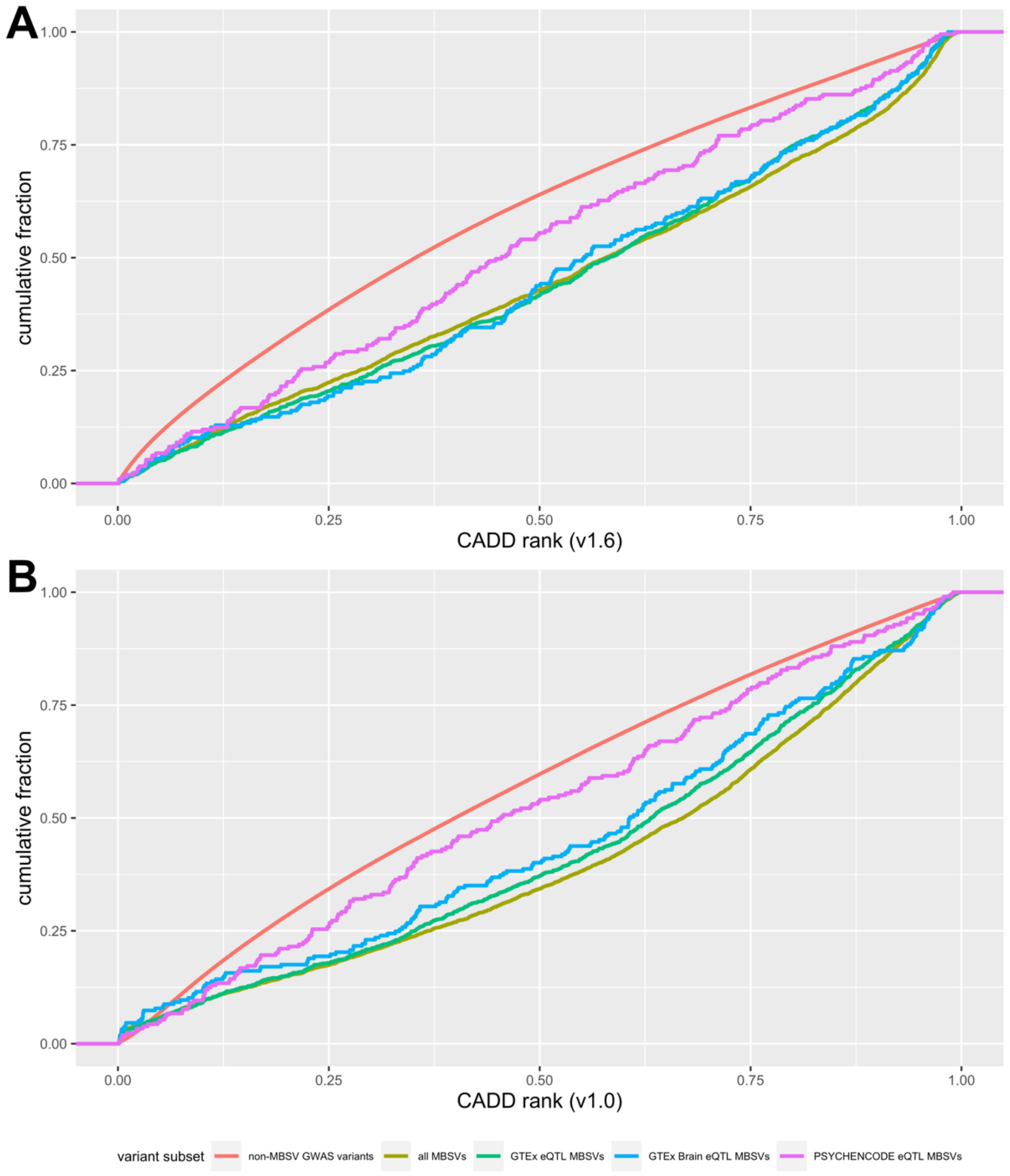


Supplementary Figure 1 | Distribution of CADD score ranks. (a, b) CADD score ranks from the latest version of the database (v1.6) (a) and the original database version (v1.0) (b) were retrieved for all MBSVs, eQTL MBSVs, and non-MBSVs. Kolmogorov-Smirnov tests were performed on each MBSV class, with non-MBSVs as the comparison group. All MBSV categories except for PSYCHENCODE eQTL MBSVs in the v1.6 database were significantly differentially distributed (Benjamini-Hochberg FDR < 0.05), with median score ranks being greater than non-MBSVs in all comparisons.

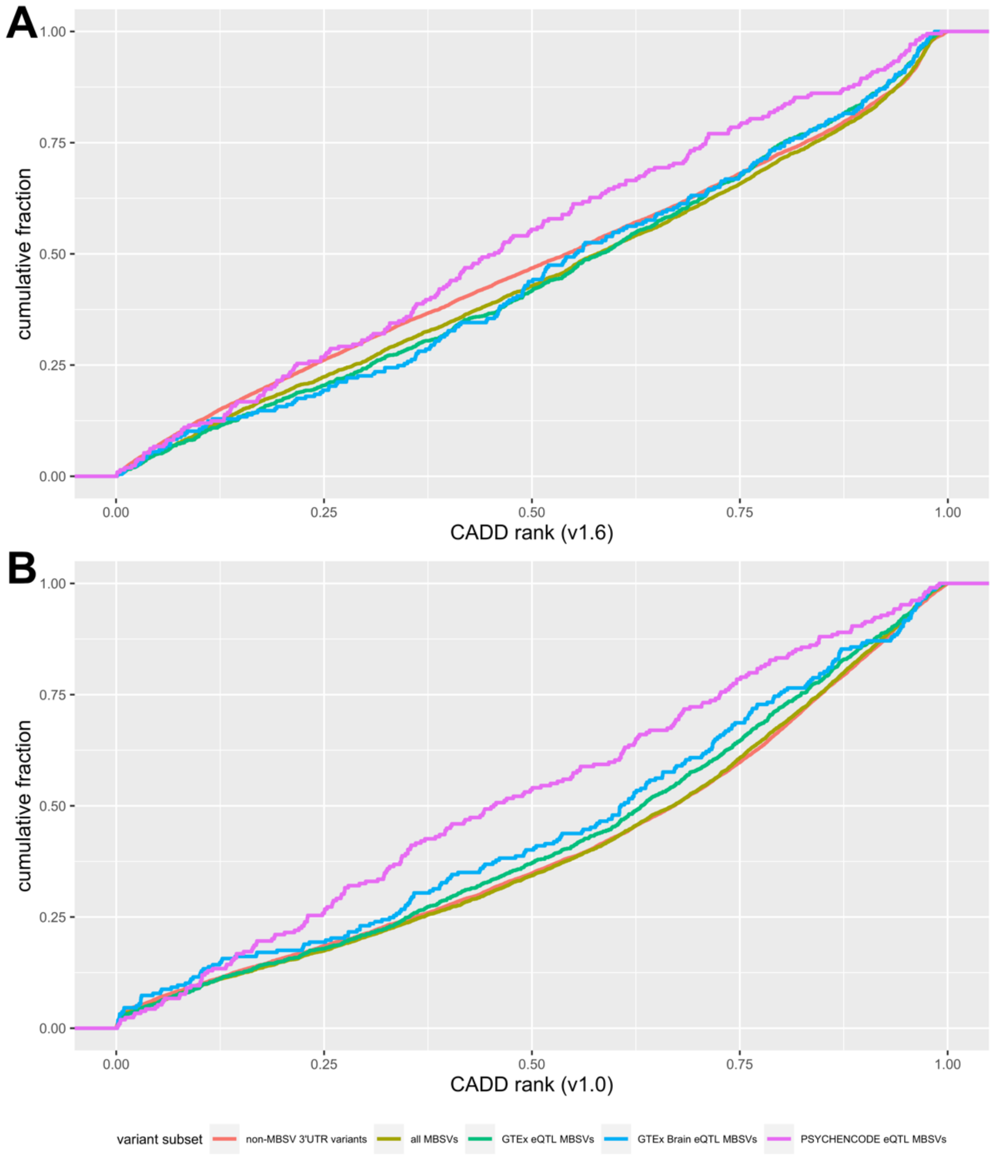


Supplementary Figure 2 | Distribution of CADD score ranks. (a, b) CADD score ranks from the latest version of the database (v1.6) (a) and the original database version (v1.0) (b) were retrieved for all MBSVs, eQTL MBSVs, and 3’ UTR-localised non-MBSVs. Kolmogorov-Smirnov tests were performed on each MBSV class, with non-MBSVs as the comparison group. All MBSV categories except for PSYCHENCODE eQTL MBSVs in the v1.6 database were significantly differentially distributed (Benjamini-Hochberg FDR < 0.05), with median score ranks being greater than 3’ UTR-localised non-MBSVs in all comparisons.

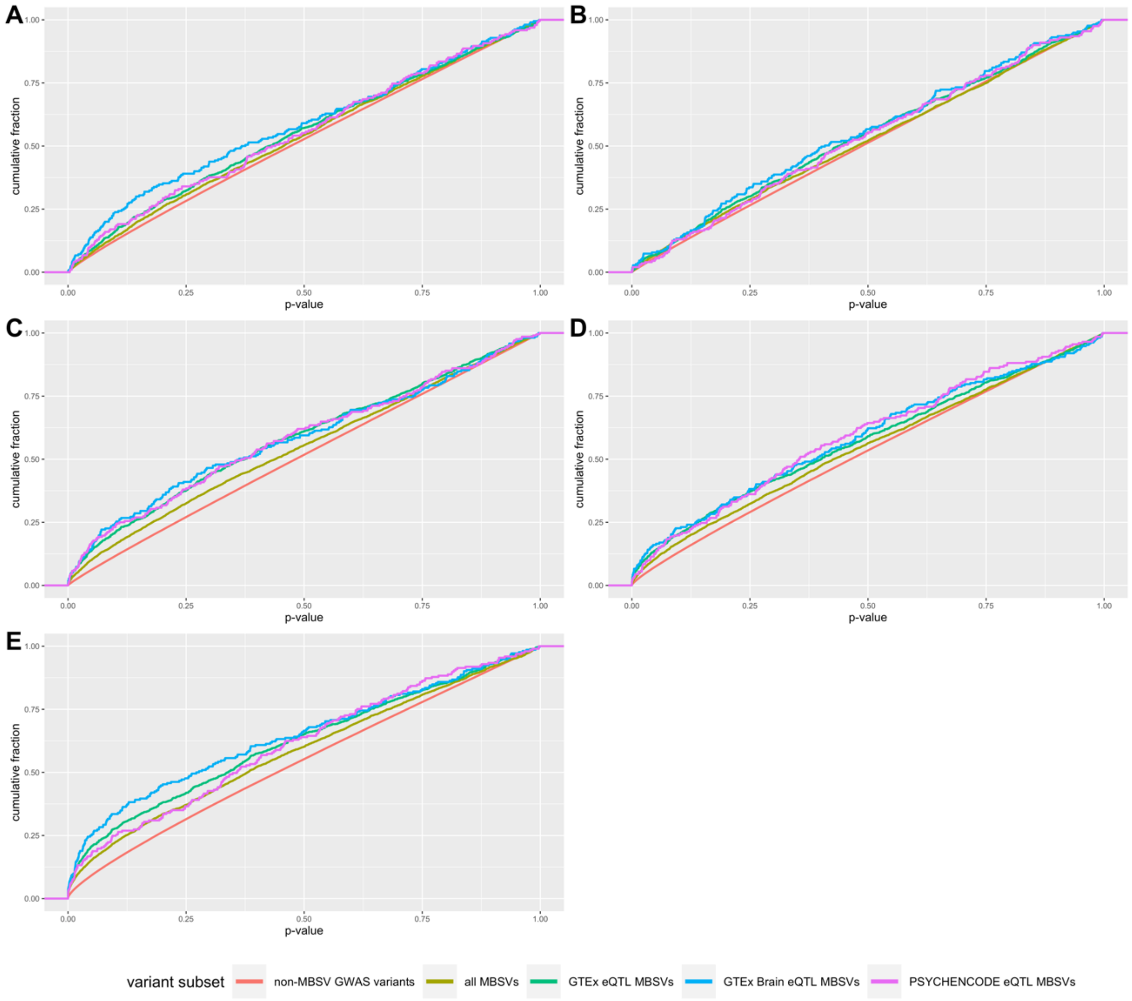


Supplementary Figure 3 | Distribution of p-values for AN, ASD, BIP, MDD, and SCZ. For each disorder, p-values for MBSVs and eQTL MBSVs were retrieved and compared to non-MBSVs with the Kolmogorov-Smirnov test. Median p-values were also compared. Significant (Benjamini-Hochberg FDR < 0.05) differences in p-value distributions of all MBSVs were identified in AN (a), ASD (b), BIP (c), MDD (d), and SCZ (e), with each of these disorders displaying a significant difference in the distribution of at least one eQTL MBSV category (GTEx eQTL MBSVs, GTEx brain eQTL MBSVs, or PSYCHENCODE eQTL MBSVs). Neither ADHD nor PTSD showed any significant differences, while OCD and TS showed a slight difference for GTEx eQTL MBSVs only (see Figure Supplementary Figure 4). In each case of altered p-value distribution, the median p-value was lower for MBSVs compared to non-MBSVs.

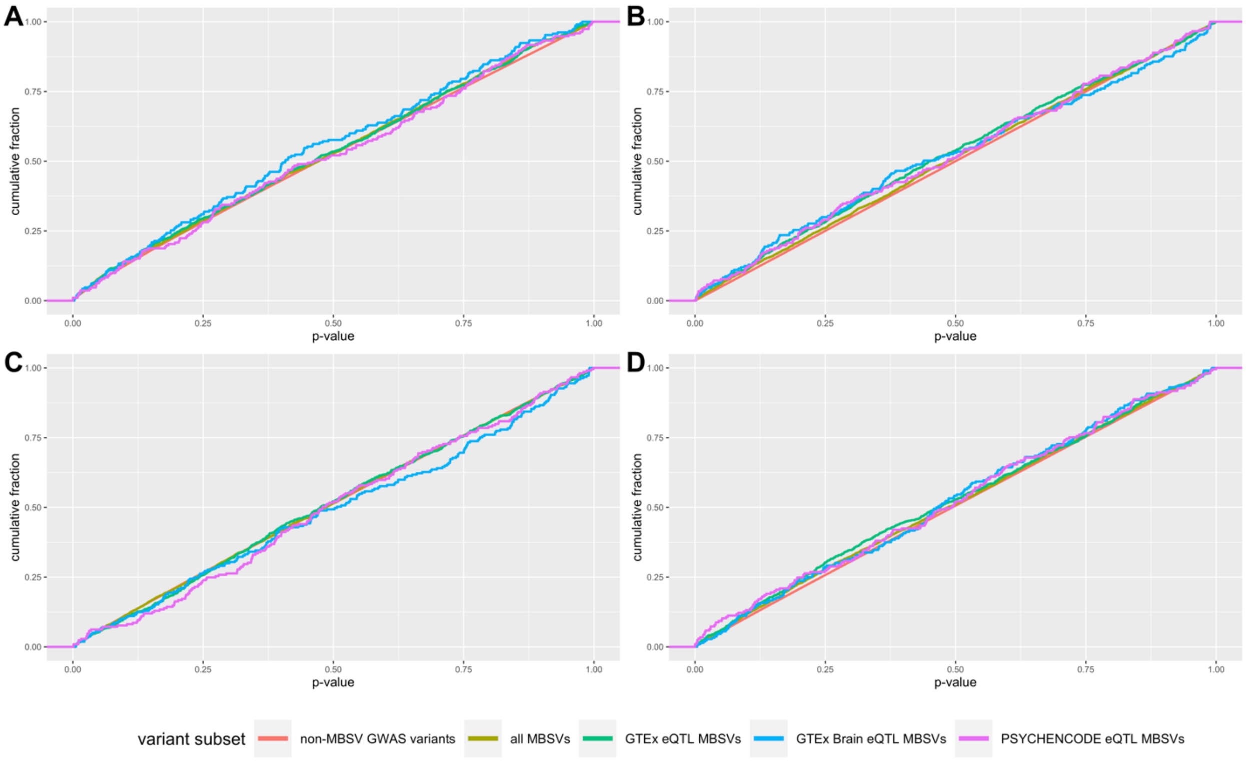


Supplementary Figure 4 | Distribution of p-values for ADHD (a), OCD (b), PTSD (c), and TS (c). For each disorder, p-values for MBSVs and eQTL MBSVs were retrieved and compared to non-MBSVs with the Kolmogorov-Smirnov test. Median p-values were also compared. No significant (Benjamini-Hochberg FDR < 0.05) differences in p-value distributions were found except for GTEx eQTL MBSVs in OCD and TS; for OCD there was a slight increase in median p-value; for TS there was a slight decrease.

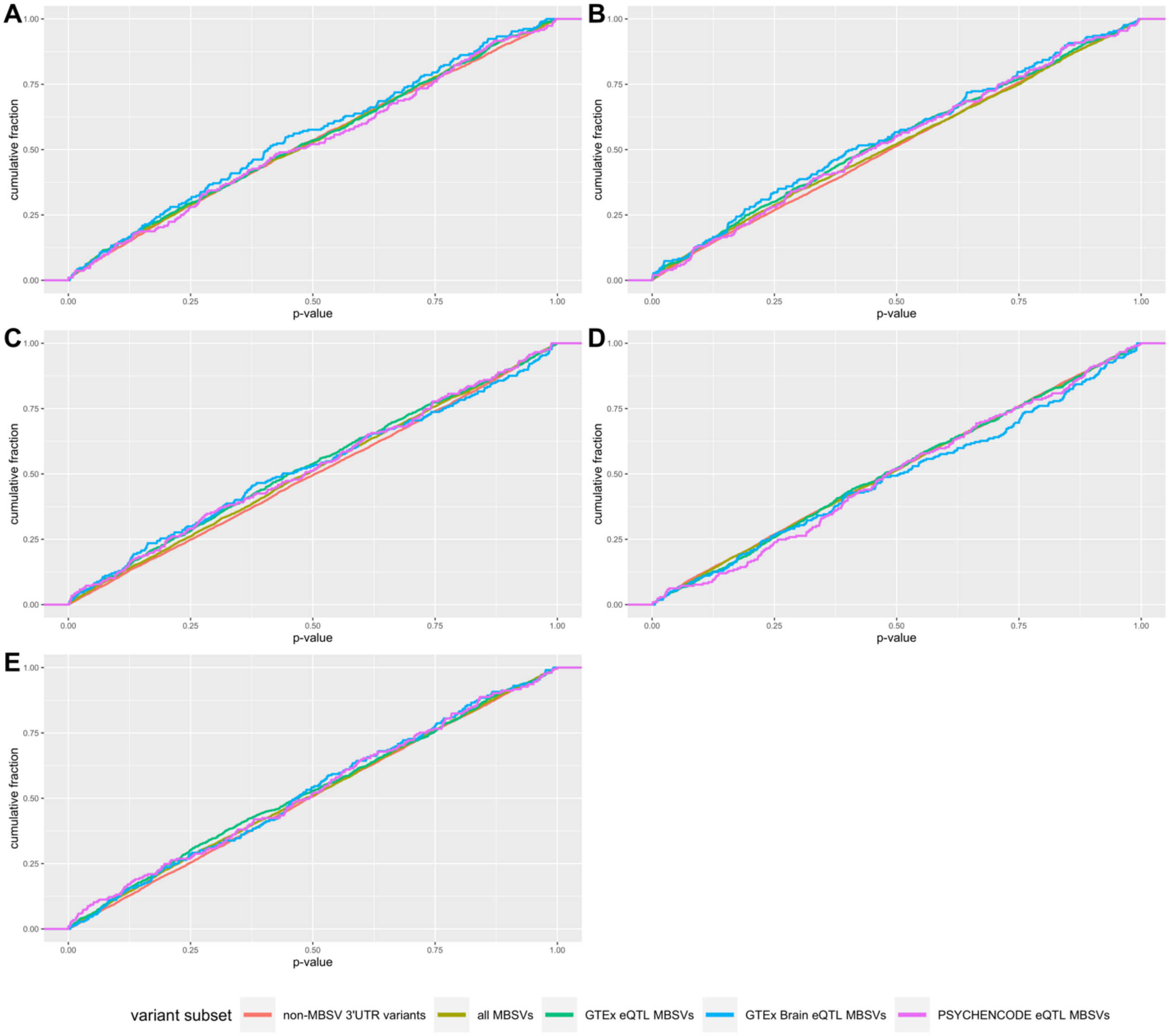


Supplementary Figure 5 | Distribution of p-values for ADHD (a), ASD (b), OCD (c), PTSD (d), and TS (e). For each disorder, p-values for MBSVs and eQTL MBSVs were retrieved and compared to 3’ UTR-localised non-MBSVs with the Kolmogorov-Smirnov test. Median p-values were also compared. No significant (Benjamini-Hochberg FDR < 0.05) differences in p-value distributions were found except for GTEx eQTL MBSVs in ASD, OCD and TS, for which the median p-values were slightly decreased.

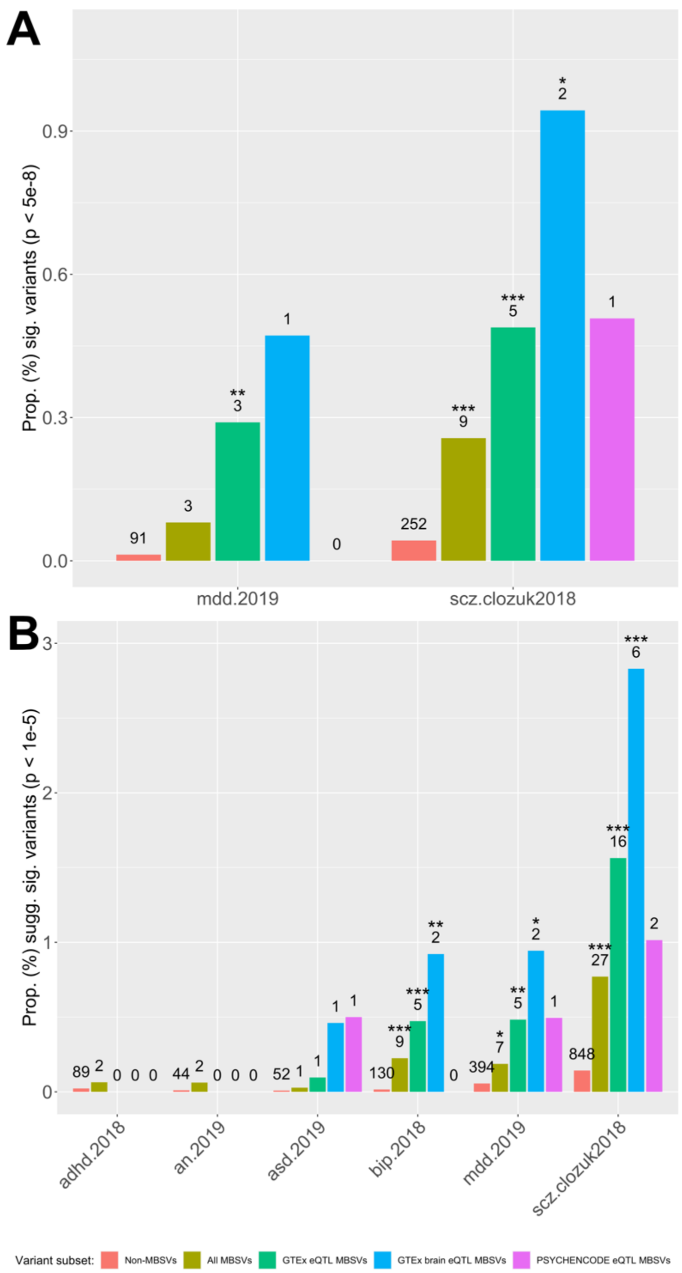


Supplementary Figure 6 | Proportions of strongly-associated variants in each disorder. (a) Variants were filtered for genome-wide significance ($p < 5x10^{-8}$). Proportions of all strongly-associated MBSVs (gold) and eQTL-annotated MBSVs (GTEx in green; GTEx brain in blue; PSYCHENCODE in pink) were compared to the proportion of strongly-associated non-MBSVs (red) using Fisher's exact test. Only disorders with at least one significant MBSV are shown. Numbers represent counts of significant variants. * = $p < 0.05$; ** = $p < 0.01$; *** = $p < 0.001$.

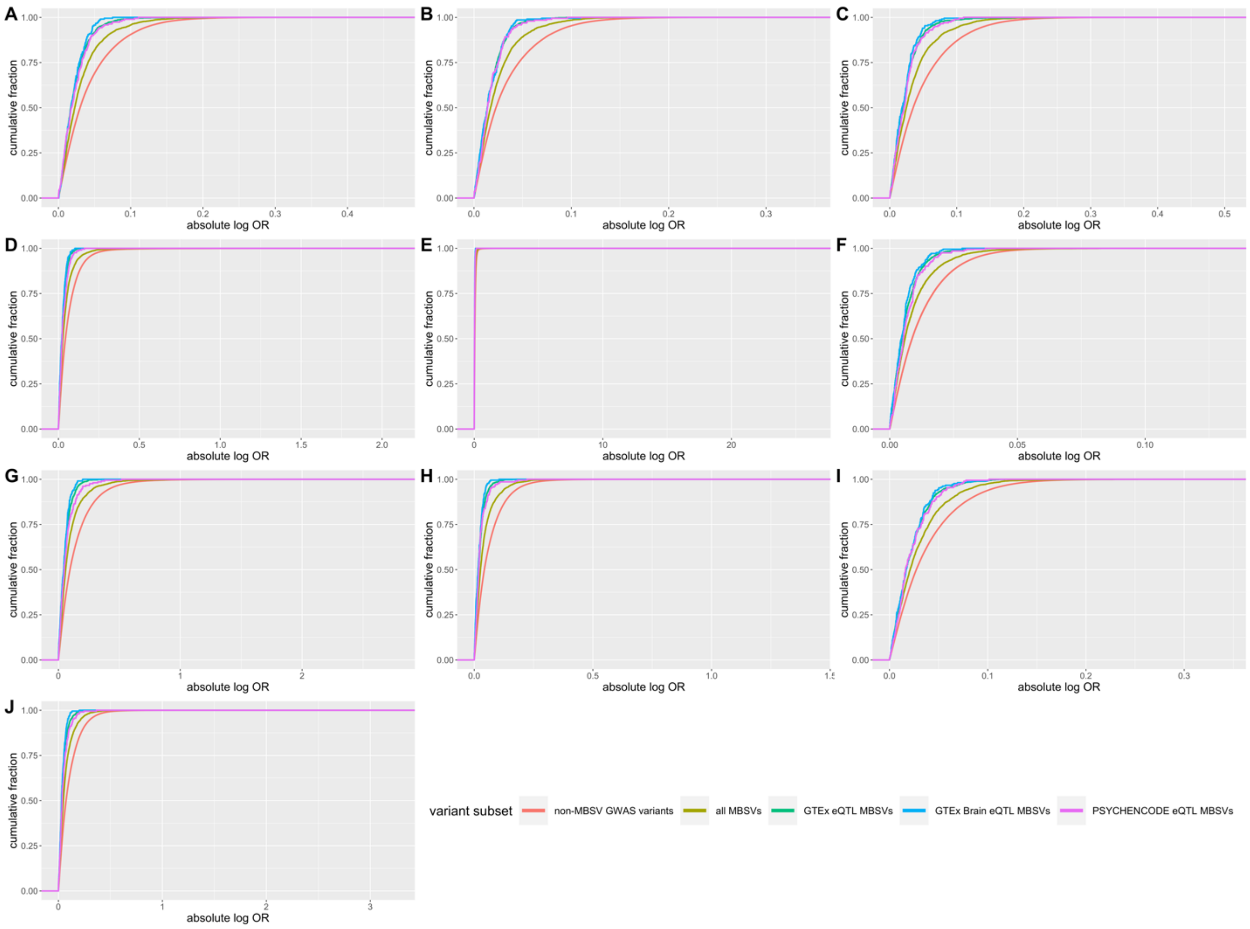


Supplementary Figure 7 | Distribution of absolute log-transformed effect sizes for MBSVs, eQTL MBSVs, and non-MBSVs for psychiatric disorders: (a) ADHD, (b) AN, (c) ASD, (d) BIP, (e) BIP, filtered for abs(logOR) < 2.1, (f) MDD, (g) OCD, (h) PTSD, (i) SCZ, (j) TS. Kolmogorov-Smirnov tests were used to compare MBSV classes to non-MBSVs. Median effect sizes were compared. In each case, MBSVs were significantly enriched for smaller effect sizes compared to non-MBSVs.

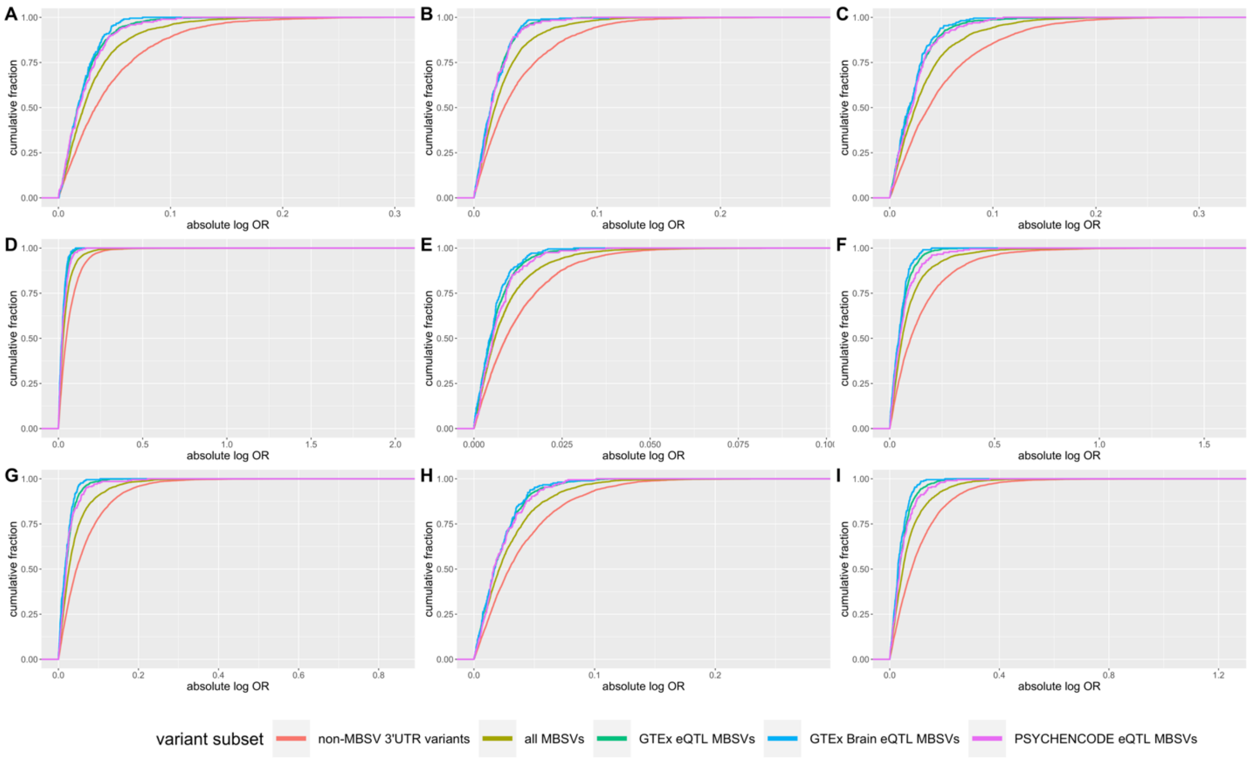


Supplementary Figure 8 | Distribution of absolute log-transformed effect sizes for MBSVs, eQTL MBSVs, and 3' UTR-localised non-MBSVs for psychiatric disorders: (a) ADHD, (b) AN, (c) ASD, (d) BIP, (e) MDD, (f) OCD, (g) PTSD, (h) SCZ, (i) TS. Kolmogorov-Smirnov tests were used to compare MBSV classes to non-MBSVs. Median effect sizes were compared. In each case, MBSVs were significantly enriched for smaller effect sizes compared to 3' UTR-localised non-MBSVs.

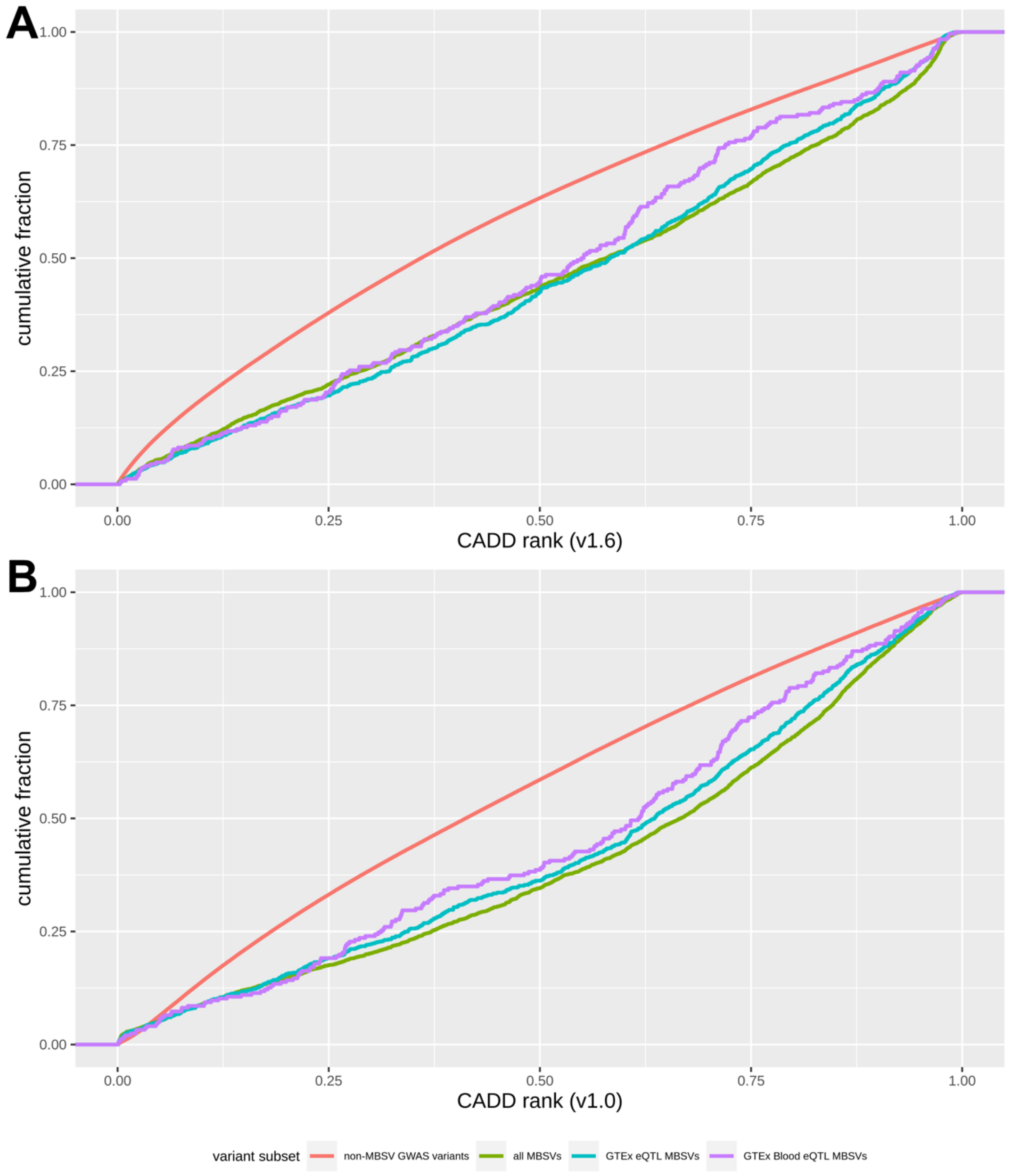


Supplementary Figure 9 | Distribution of CADD score ranks for variants tested in nonpsychiatric traits, with MBSVs determined relative to blood gene and miRNA expression. (a, b) CADD score ranks from the latest version of the database (v1.6) (a) and the original database version (v1.0) (b) were retrieved for all MBSVs, eQTL MBSVs, and non-MBSVs. Kolmogorov-Smirnov tests were performed on each MBSV class, with non-MBSVs as the comparison group.

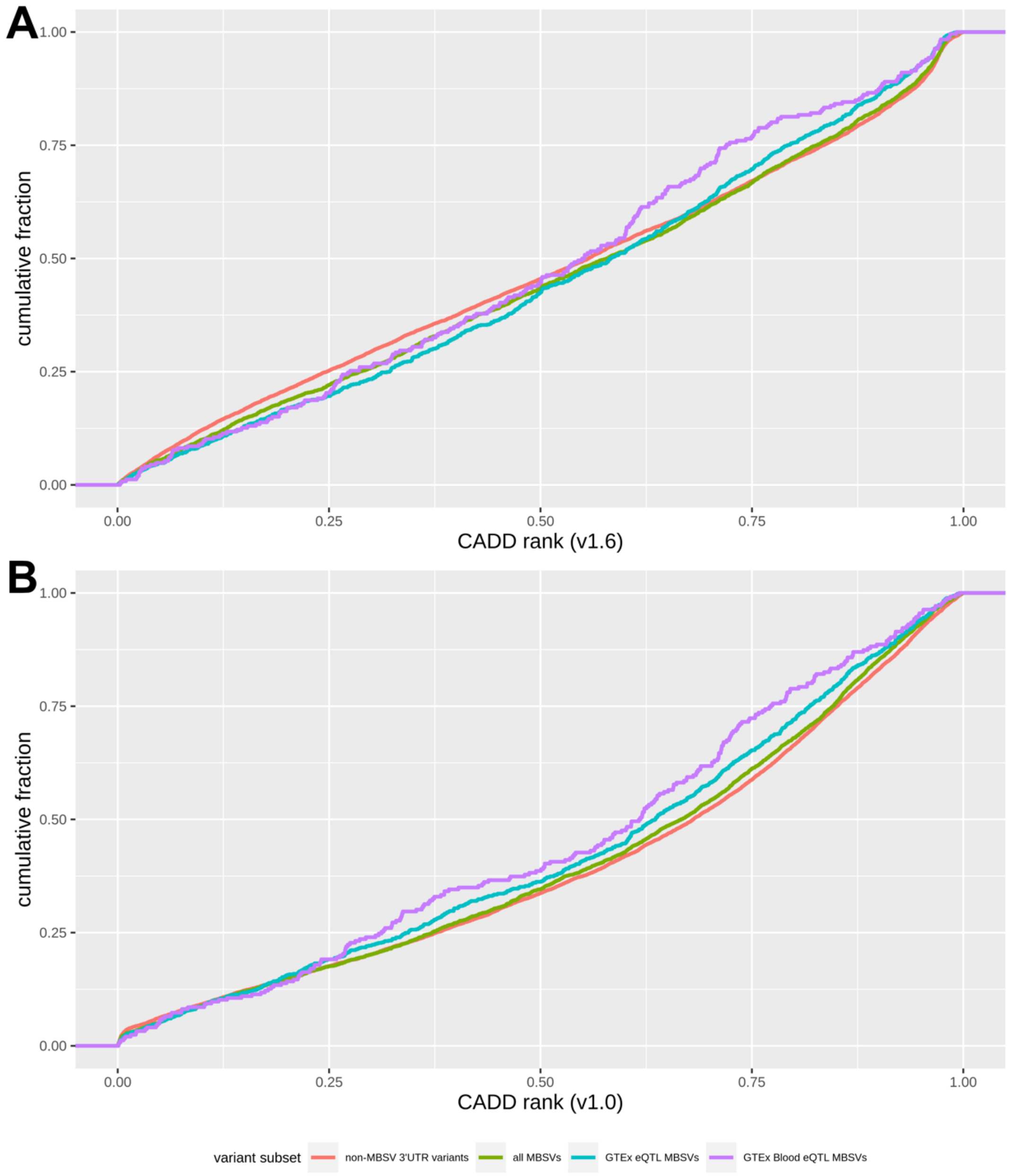


Supplementary Figure 10 | Distribution of CADD score ranks for variants tested in nonpsychiatric traits, with MBSVs determined relative to blood gene and miRNA expression. (a, b) CADD score ranks from the latest version of the database (v1.6) (a) and the original database version (v1.0) (b) were retrieved for all MBSVs, eQTL MBSVs, and 3’ UTR-localised non-MBSVs. Kolmogorov-Smirnov tests were performed on each MBSV class, with non-MBSVs as the comparison group.

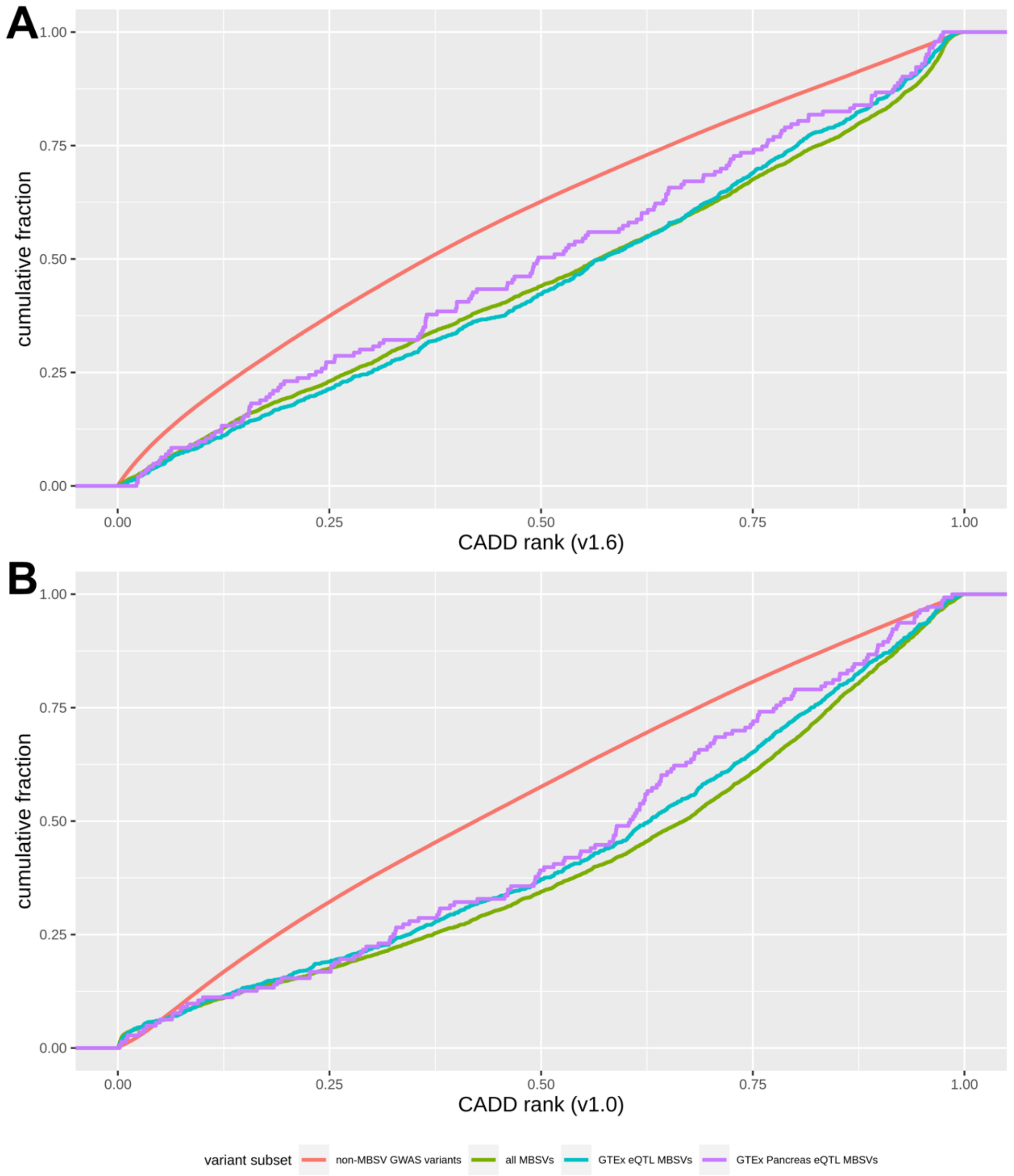


Supplementary Figure 11 | Distribution of CADD score ranks for variants tested in type 2 diabetes, with MBSVs determined relative to pancreas gene and miRNA expression. (a, b) CADD score ranks from the latest version of the database (v1.6) (a) and the original database version (v1.0) (b) were retrieved for all MBSVs, eQTL MBSVs, and non-MBSVs. Kolmogorov-Smirnov tests were performed on each MBSV class, with non-MBSVs as the comparison group.

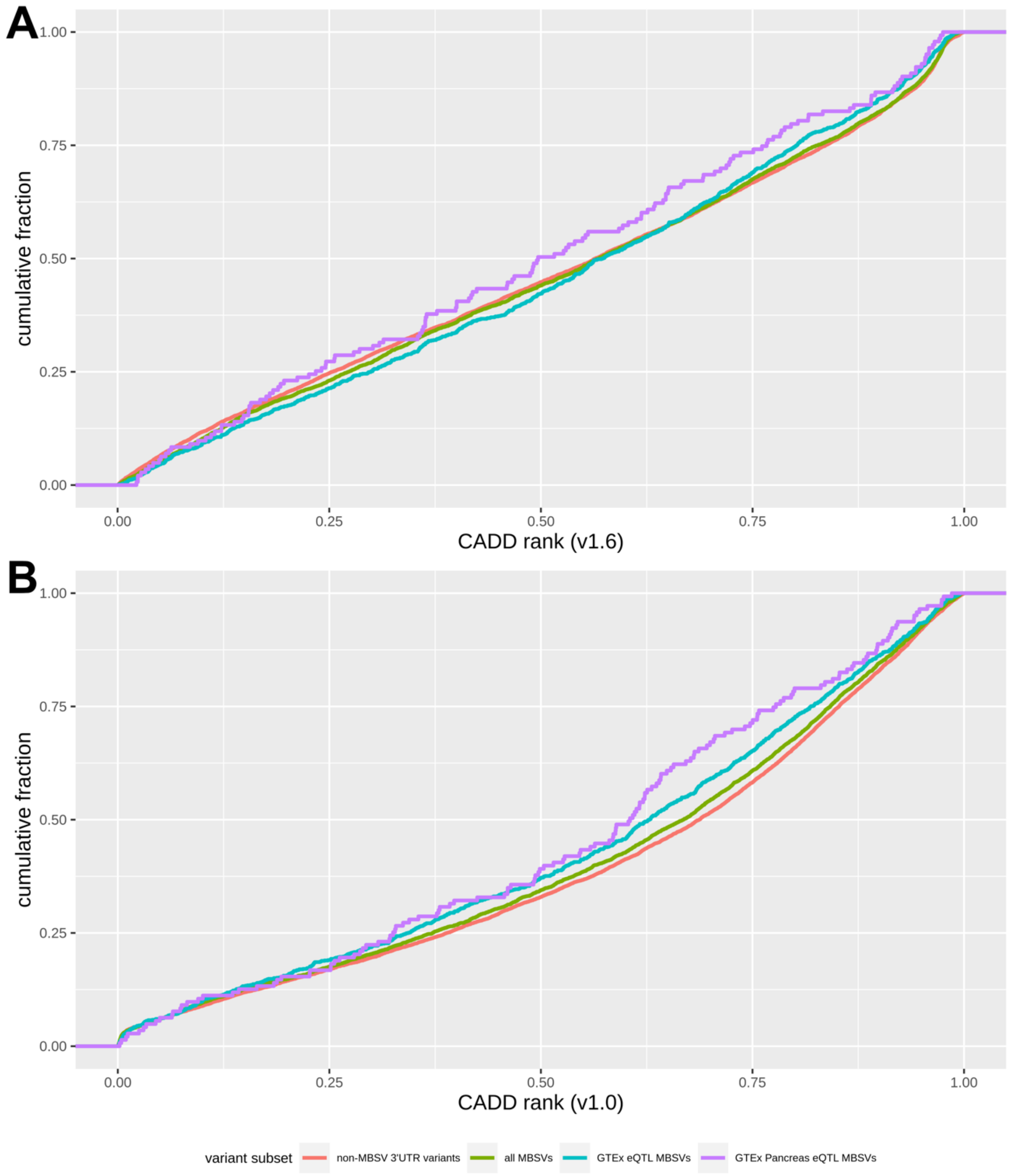


Supplementary Figure 12 | Distribution of CADD score ranks for variants tested in type 2 diabetes, with MBSVs determined relative to pancreas gene and miRNA expression. (a, b) CADD score ranks from the latest version of the database (v1.6) (a) and the original database version (v1.0) (b) were retrieved for all MBSVs, eQTL MBSVs, and 3' UTR-localised non-MBSVs. Kolmogorov-Smirnov tests were performed on each MBSV class, with non-MBSVs as the comparison group.

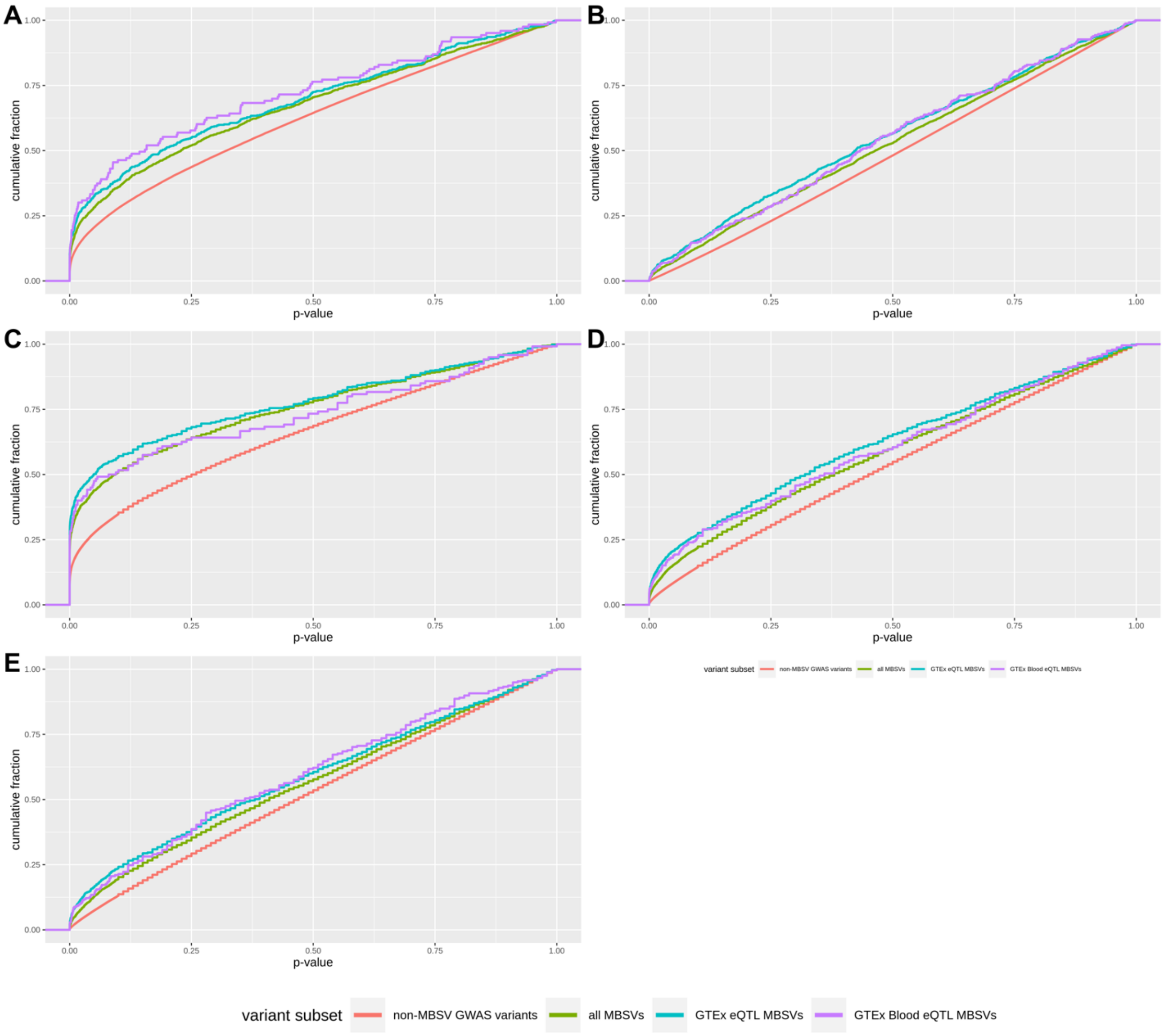


Supplementary Figure 13 | Distribution of p-values for BMI (a), CAD (b), height (c), T2D (d) and T2D (BMI adj.) (e), with MBSVs determined relative to blood gene and miRNA expression. For each disorder, p-values for MBSVs and eQTL MBSVs were retrieved and compared to non-MBSVs with the Kolmogorov-Smirnov test.

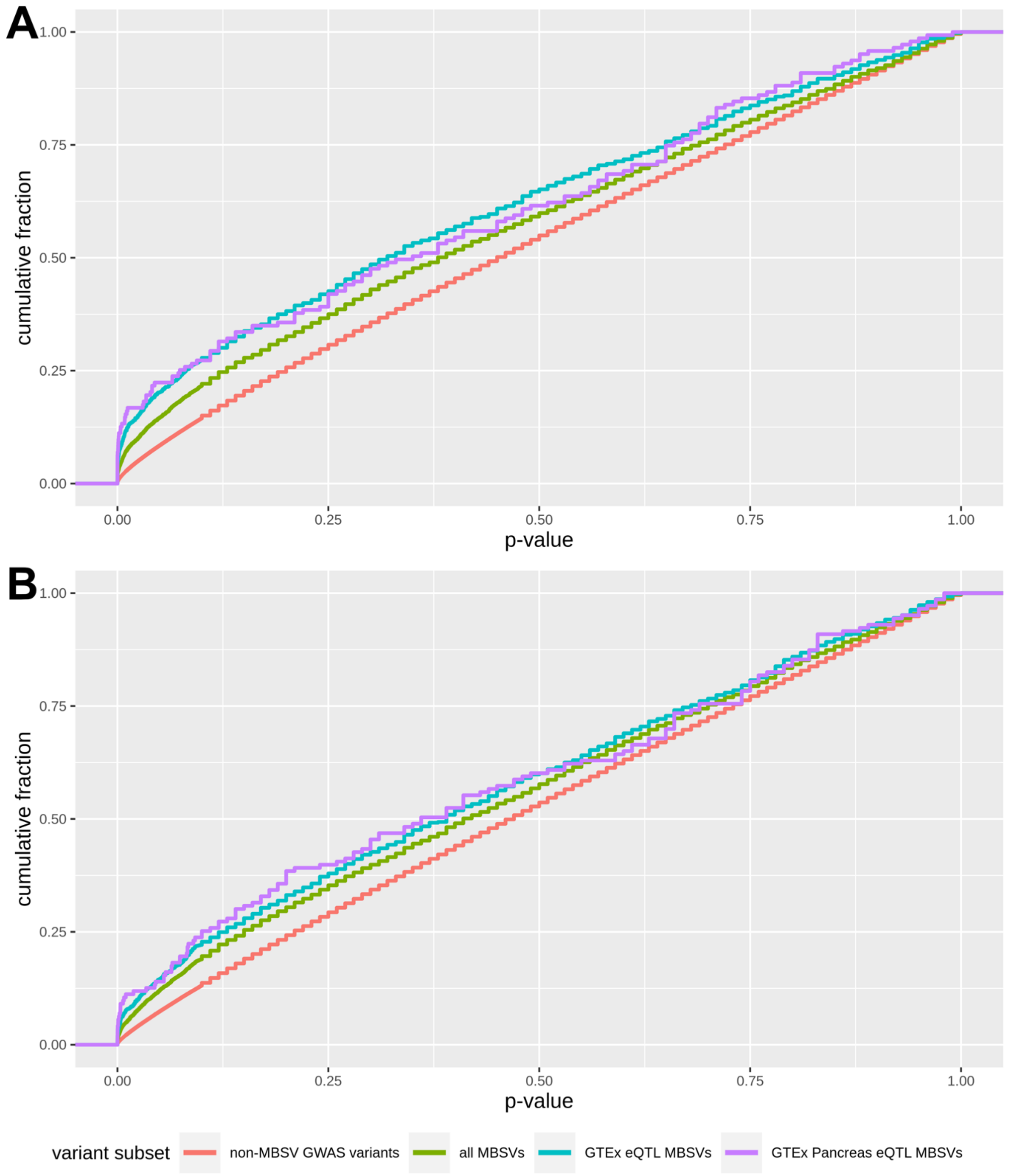


Supplementary Figure 14 | Distribution of p-values for T2D (a) and T2D (BMI adj.) (b), with MBSVs determined relative to pancreas gene and miRNA expression. For each disorder, p-values for MBSVs and eQTL MBSVs were retrieved and compared to non-MBSVs with the Kolmogorov-Smirnov test.

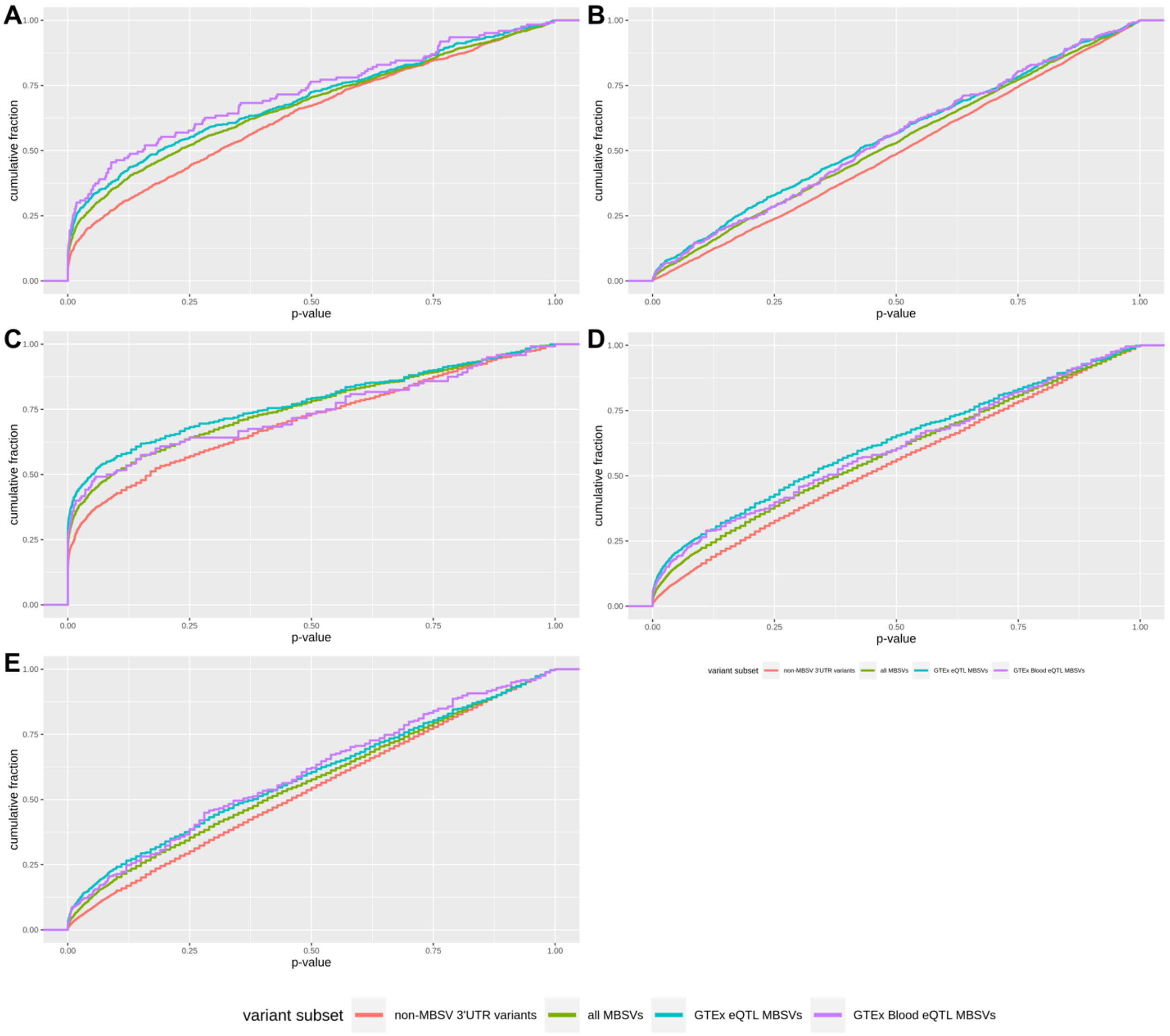


Supplementary Figure 15 | Distribution of p-values for BMI (a), CAD (b), height (c), T2D (d) and T2D (BMI adj.) (e), with MBSVs determined relative to blood gene and miRNA expression. For each disorder, p-values for MBSVs and eQTL MBSVs were retrieved and compared to 3' UTR-localised non-MBSVs with the Kolmogorov-Smirnov test.

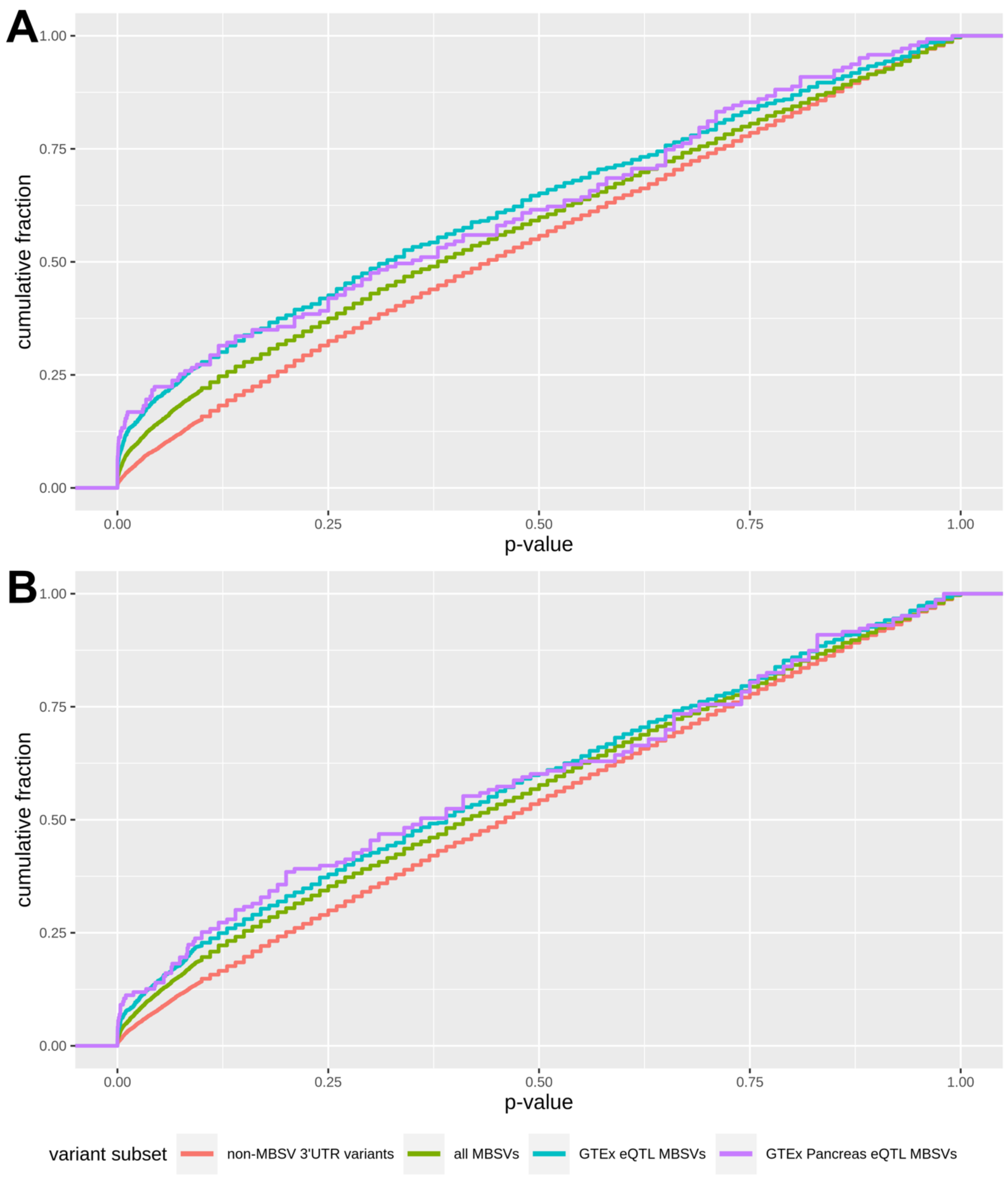


Supplementary Figure 16 | Distribution of p-values for T2D (a) and T2D (BMI adj.) (b), with MBSVs determined relative to pancreas gene and miRNA expression. For each disorder, p-values for MBSVs and eQTL MBSVs were retrieved and compared to 3’ UTR-localised non-MBSVs with the Kolmogorov-Smirnov test.

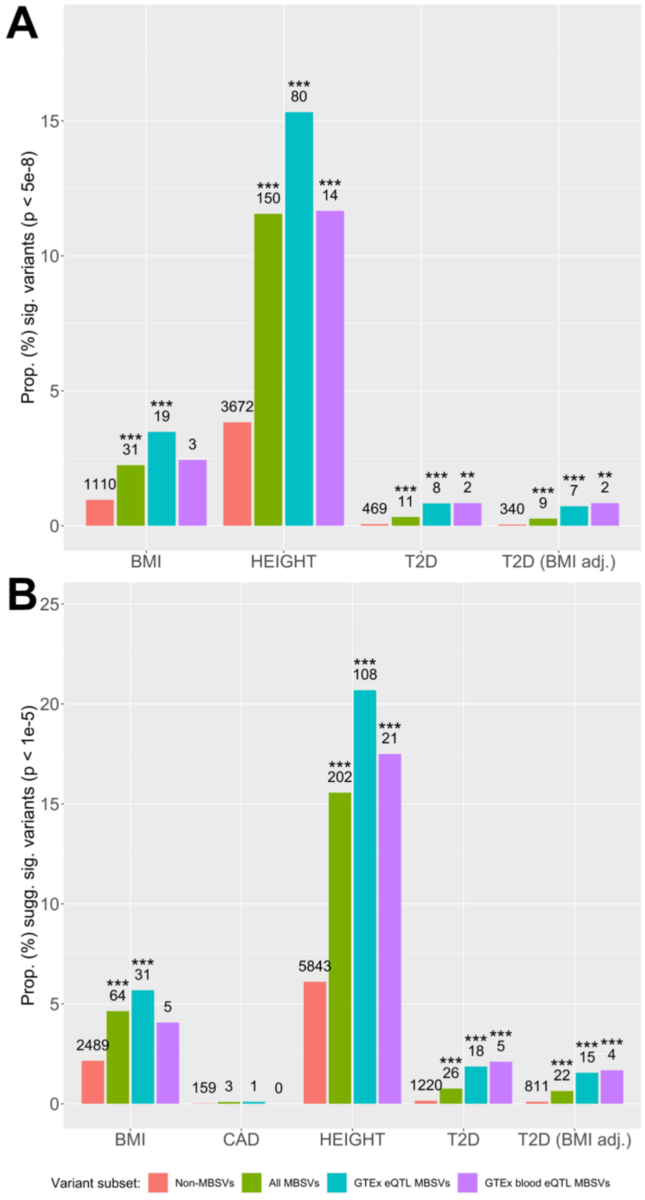


Supplementary Figure 17 | Proportions of strongly-associated variants in each non-psychiatric trait, with MBSVs determined relative to blood gene and miRNA expression. Variants were filtered for genome-wide significance ($p < 5x10^{-8}$). Proportions of all strongly-associated MBSVs (green) and eQTL-annotated MBSVs (GTEx in blue; GTEx blood in purple) were compared to the proportion of strongly-associated non-MBSVs (red) using Fisher's exact test. Only traits with at least one significant MBSV are shown. Numbers represent counts of significant variants. * = p < 0.05; ** = p < 0.01; *** = p < 0.001.

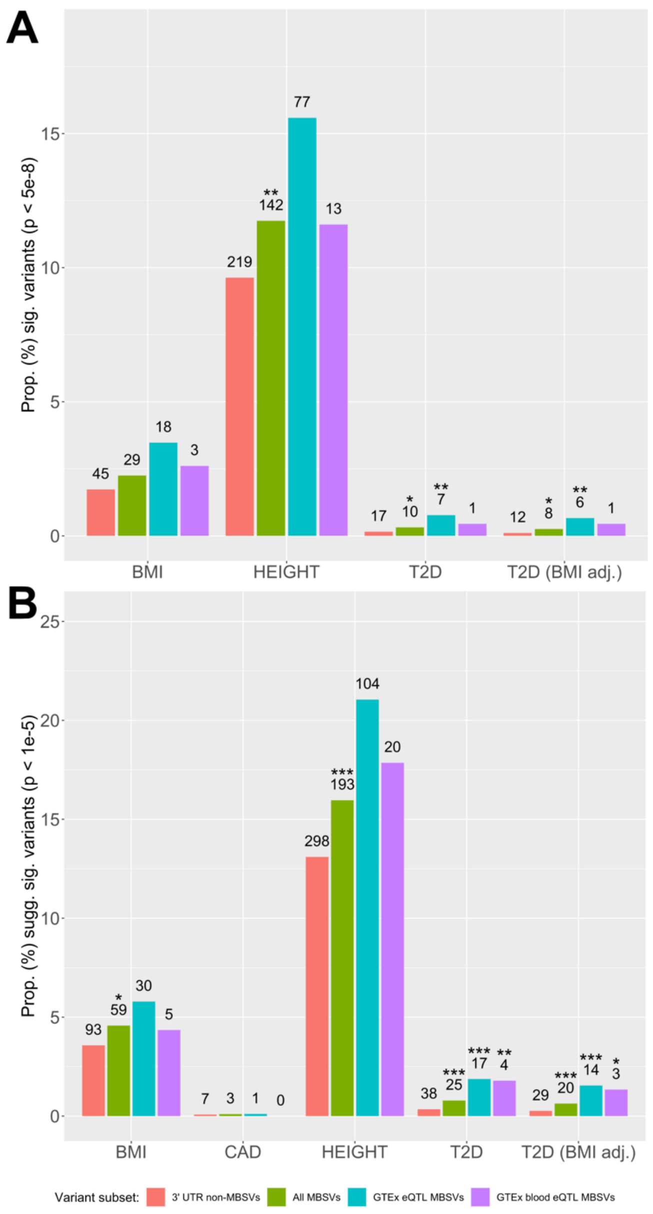


Supplementary Figure 18 | Proportions of strongly-associated variants in each non-psychiatric trait, with MBSVs determined relative to blood gene and miRNA expression. Variants were filtered for genome-wide significance ($p < 5 \times 10^{-8}$). Proportions of all strongly-associated MBSVs (green) and eQTL-annotated MBSVs (GTEx in blue; GTEx blood in purple) were compared to the proportion of strongly-associated 3'UTR-localised non-MBSVs (red) using Fisher's exact test. Only traits with at least one significant MBSV are shown. Numbers represent counts of significant variants. * = $p < 0.05$; ** = $p < 0.01$; *** = $p < 0.001$.

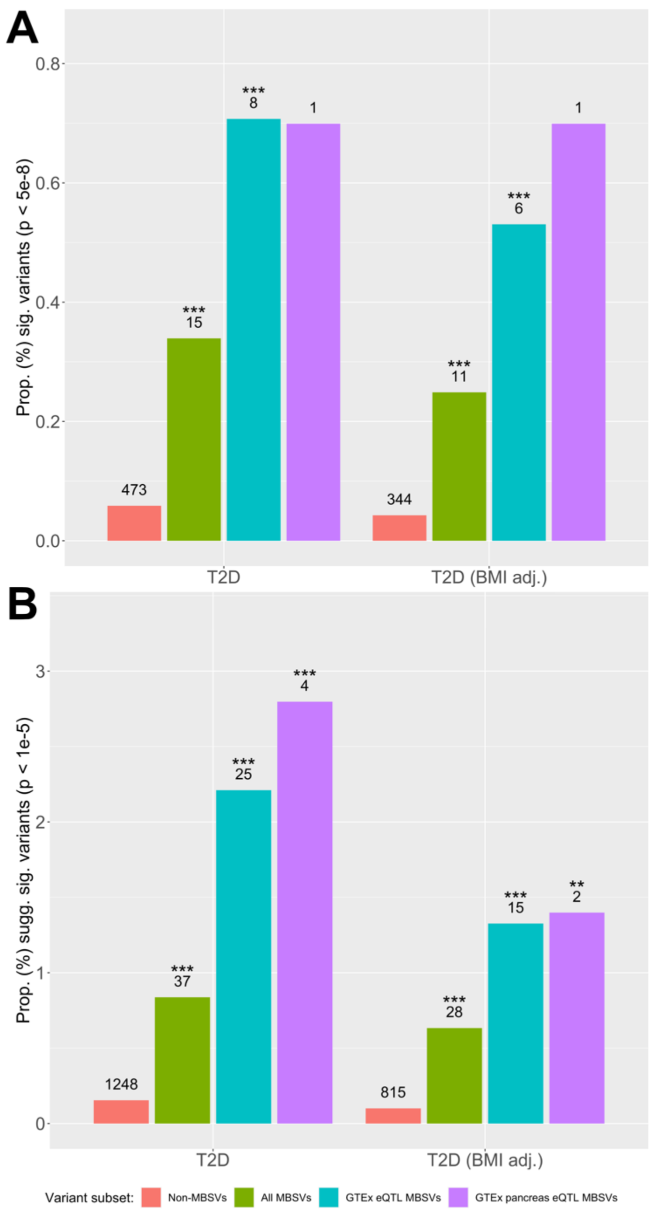


Supplementary Figure 19 | Proportions of strongly-associated variants in T2D and T2D (BMI adj.), with MBSVs determined relative to pancreas gene and miRNA expression. Variants were filtered for genome-wide significance ($p < 5x10^{-8}$). Proportions of all strongly-associated MBSVs (green) and eQTL-annotated MBSVs (GTEx in blue; GTEx blood in purple) were compared to the proportion of strongly-associated non-MBSVs (red) using Fisher's exact test. Only traits with at least one significant MBSV are shown. Numbers represent counts of significant variants. * = p < 0.05; ** = p < 0.01; *** = p < 0.001.

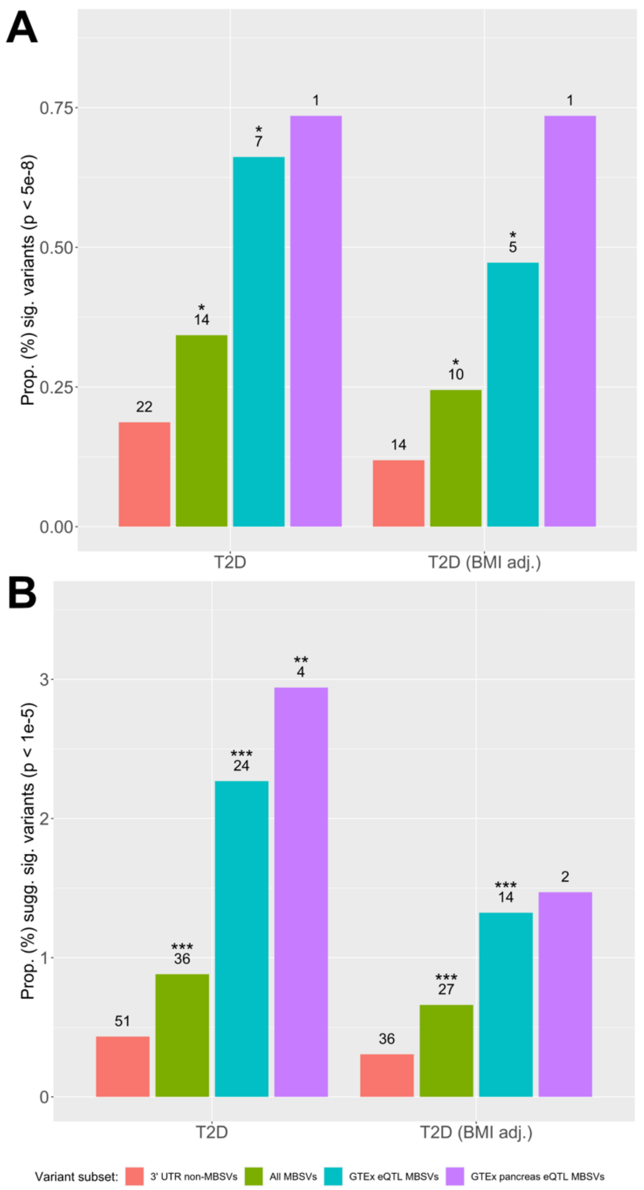


Supplementary Figure 20 | Proportions of strongly-associated variants in T2D and T2D (BMI adj.), with MBSVs determined relative to pancreas gene and miRNA expression. Variants were filtered for genome-wide significance ($p < 5x10^{-8}$). Proportions of all strongly-associated MBSVs (green) and eQTL-annotated MBSVs (GTEx in blue; GTEx blood in purple) were compared to the proportion of strongly-associated 3'UTR-localised non-MBSVs (red) using Fisher's exact test. Only traits with at least one significant MBSV are shown. Numbers represent counts of significant variants. * = p < 0.05; ** = p < 0.01; *** = p < 0.001.

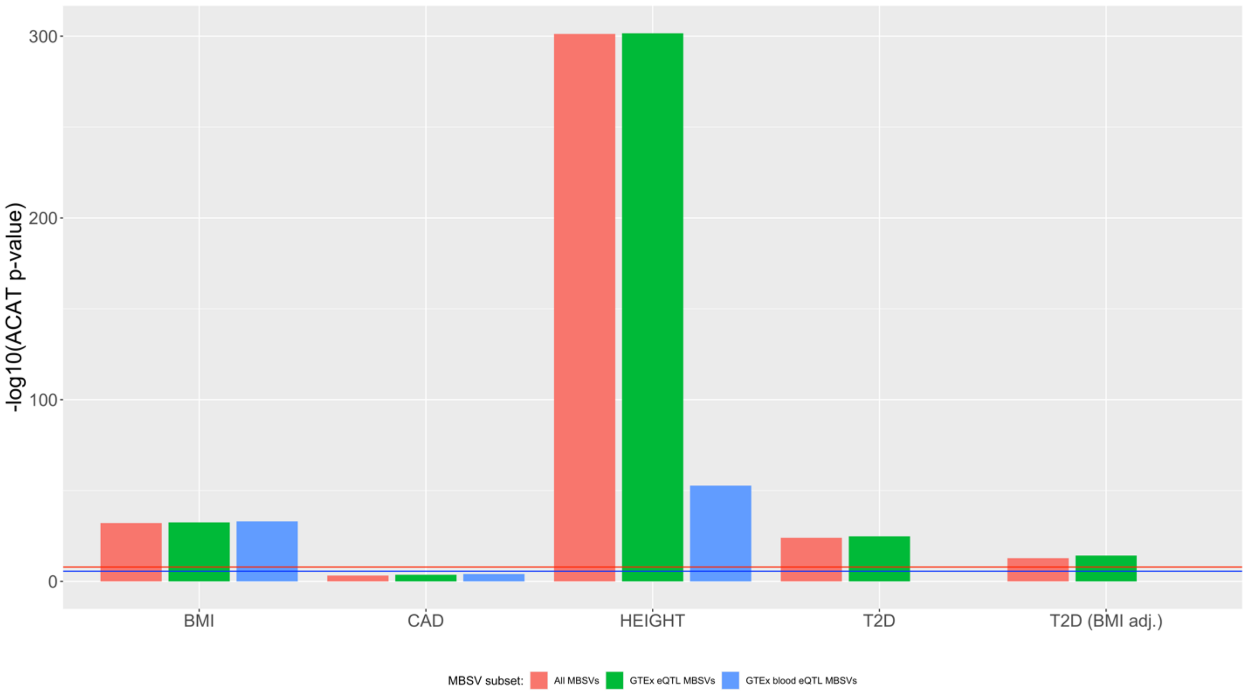


Supplementary Figure 21 | Aggregated p-values for MBSVs in each non-psychiatric trait, with MBSVs determined relative to blood gene and miRNA expression. P-values for all MBSVs (red) and eQTL-annotated MBSVs (GTEx in green; GTEx blood in blue) were aggregated using the ACAT method into one test statistic per disorder. Red line = genome-wide significance corrected for multiple tests ($1.25x10^{-8}$); blue = liberal significance threshold corrected for multiple tests ($2.5x10^{-6}$).

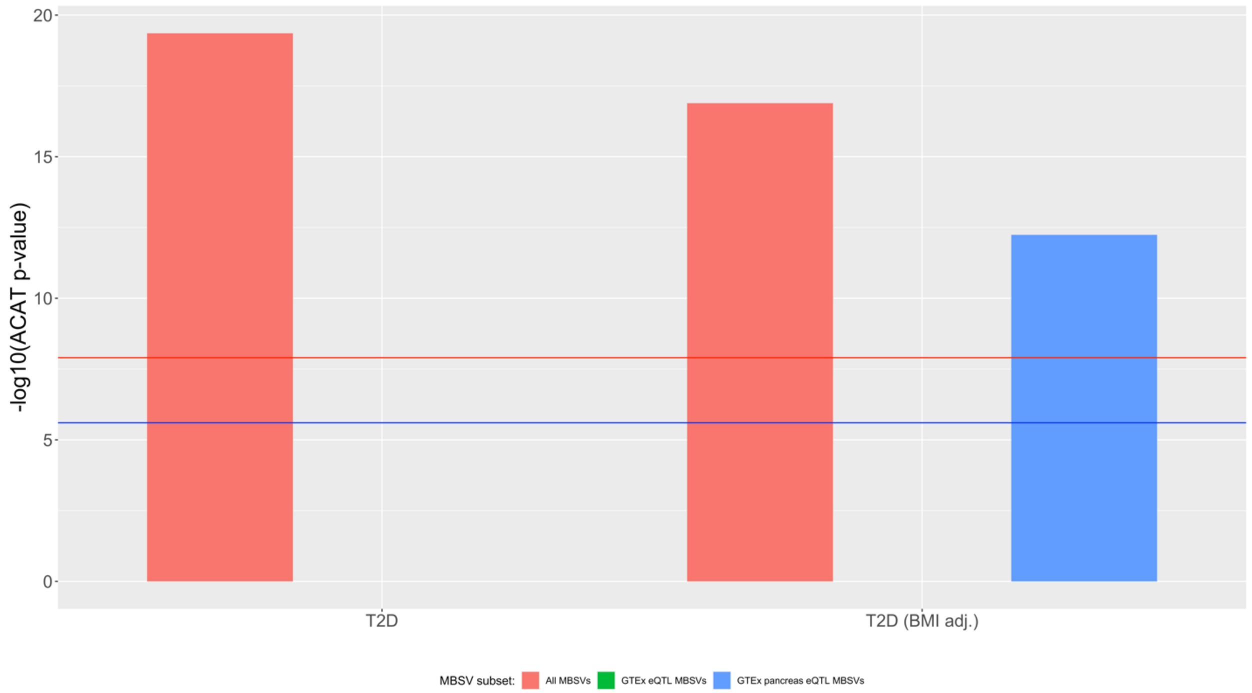


Supplementary Figure 22 | Aggregated p-values for MBSVs in T2D and T2D (BMI adj.), with MBSVs determined relative to pancreas gene and miRNA expression. P-values for all MBSVs (red) and eQTL-annotated MBSVs (GTEx in green; GTEx blood in blue) were aggregated using the ACAT method into one test statistic per disorder. Red line = genome-wide significance corrected for multiple tests ($1.25 \times 10^{-8}$); blue = liberal significance threshold corrected for multiple tests ($2.5 \times 10^{-6}$).

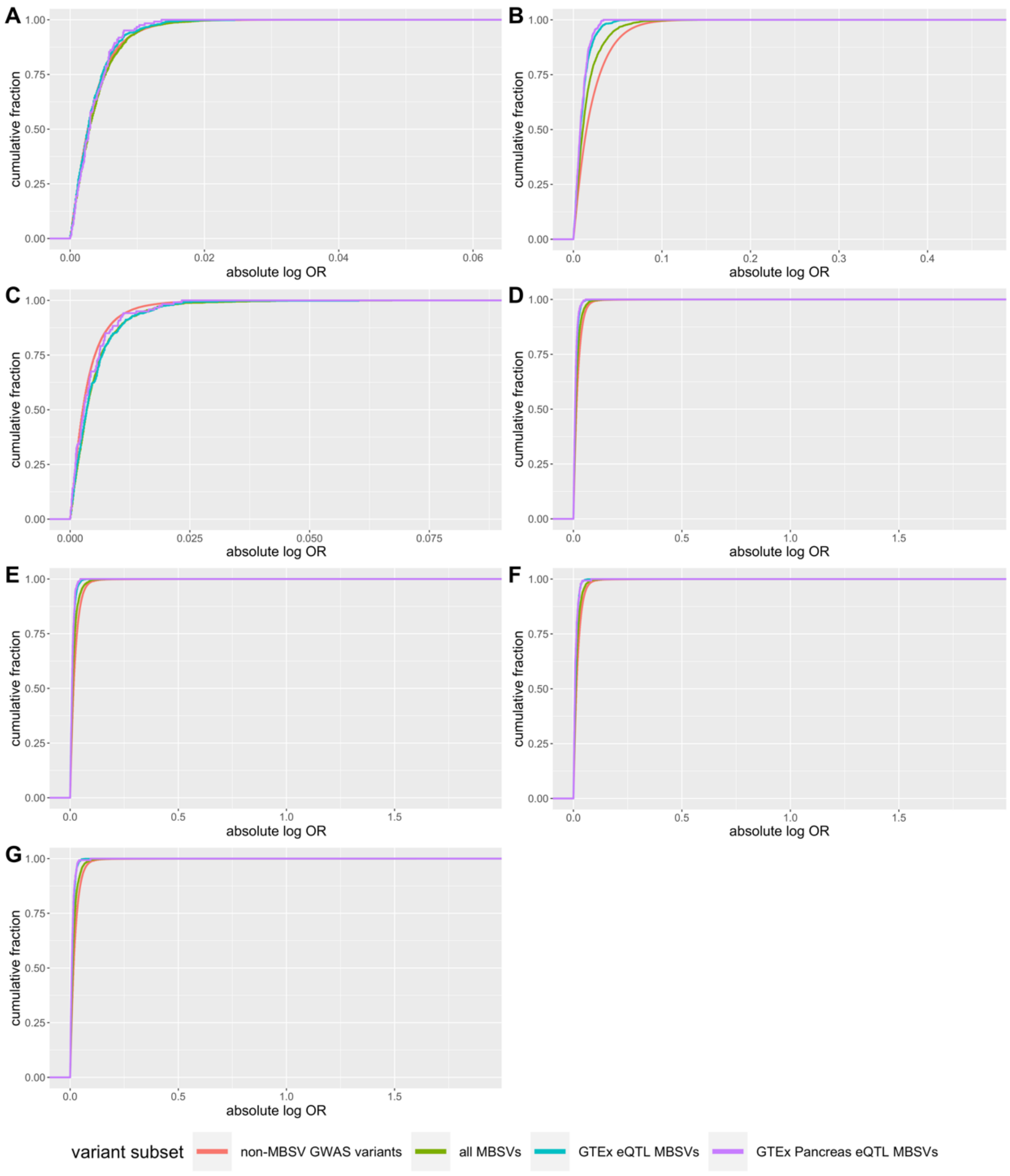


Supplementary Figure 23 | Distribution of absolute log-transformed effect sizes for MBSVs, eQTL MBSVs, and non-MBSVs for non-psychiatric traits: (a) BMI, (b) CAD, (c) height, (d) T2D (blood), (e) T2D (BMI adj., blood), (f) T2D (pancreas), (g) T2D (BMI adj., pancreas). Kolmogorov-Smirnov tests were used to compare MBSV classes to non-MBSVs. Median effect sizes were compared. In each case, MBSVs were significantly enriched for smaller effect sizes compared to non-MBSVs.

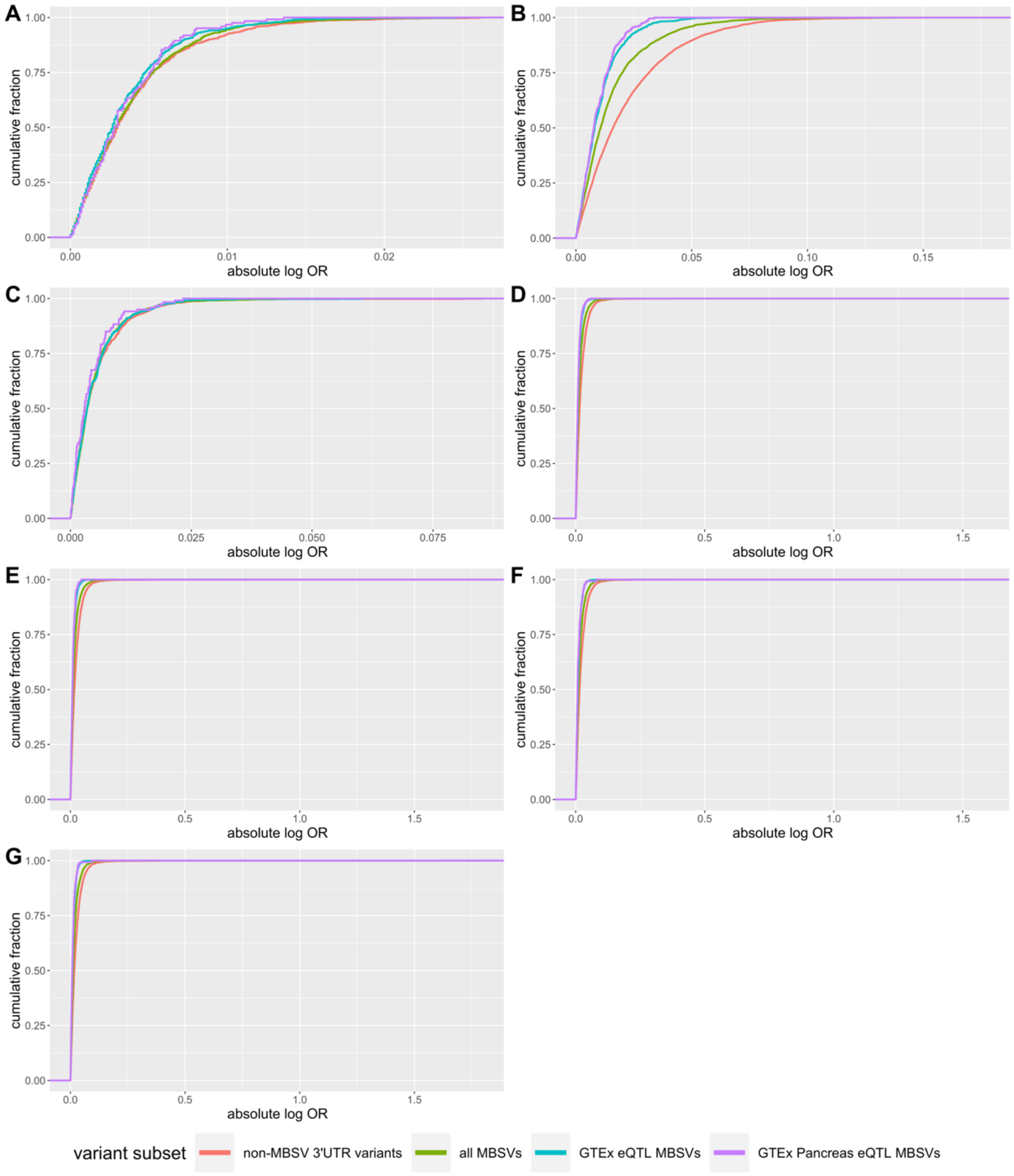


Supplementary Figure 24 | Distribution of absolute log-transformed effect sizes for MBSVs, eQTL MBSVs, and 3' UTR-localised non-MBSVs for non-psychiatric traits: (a) BMI, (b) CAD, (c) height, (d) T2D (blood), (e) T2D (BMI adj., blood), (f) T2D (pancreas), (g) T2D (BMI adj., pancreas). Kolmogorov-Smirnov tests were used to compare MBSV classes to non-MBSVs. Median effect sizes were compared. In each case, MBSVs were significantly enriched for smaller effect sizes compared to 3' UTR-localised non-MBSVs.
